# Tumor-Intrinsic Response to IFNγ Shapes the Tumor Microenvironment and Anti-PD-1 Response in NSCLC

**DOI:** 10.1101/531236

**Authors:** Bonnie L. Bullock, Abigail K. Kimball, Joanna M. Poczobutt, Howard Y. Li, Jeff W. Kwak, Alexander J. Neuwelt, Amber M. Johnson, Emily Kleczko, Rachael Kaspar, Katharina Hopp, Erin Schenk, Mary C. M. Weiser-Evans, Eric T. Clambey, Raphael A. Nemenoff

## Abstract

Targeting PD-1/ PD-L1 is only effective in ~20% of lung cancer patients, but determinants of this response are poorly defined. We previously observed differential responses of two murine K-Ras lung cancer cell lines to anti-PD-1 therapy: CMT167 tumors were eliminated while LLC tumors were resistant. The goal of this study was to define mechanism(s) mediating this difference. RNA-Seq analysis of cancer cells recovered from lung tumors revealed that CMT167 cells induced an IFNγ signature that was absent in LLC cells. Silencing *Ifngr1* in CMT167 resulted in tumors resistant to IFNγ and anti-PD-1 therapy. Conversely, LLC cells had high basal expression of *Socs1*, an inhibitor of IFNγ. Silencing *Socs1* increased response to IFNγ *in vitro* and sensitized tumors to anti-PD-1. This was associated with a reshaped TME, characterized by enhanced T cell infiltration and enrichment of PD-L1 high myeloid cells. These studies demonstrate that targeted enhancement of tumor-intrinsic IFNγ signaling can induce of cascade of changes associated with increased therapeutic vulnerability.

**Summary:** Mechanisms regulating response to anti-PD-1 therapy in lung cancer are not well defined. This study, using orthotopic immunocompetent mouse models of lung cancer, demonstrates that intrinsic sensitivity of cancer cells to IFNγ determines anti-PD-1 responsiveness through alterations in the tumor microenvironment.

## Introduction

The development of immune checkpoint inhibitors has shown great promise in a wide variety of malignancies including lung cancer. However, only ~20% of unselected non-small cell lung cancer (NSCLC) patients respond to monotherapy targeting the PD-1/PD-L1 axis [1–3]. Previous studies have correlated multiple factors with patient response to immunotherapy. These include tumor mutational burden, the presence of neoantigens, PD-L1 expression on the surface of tumor cells and/or surrounding stromal cells, tumor-infiltrating immune cells, as well as patient smoking status [4–11]. Importantly, Ayers et. al defined an interferon-gamma (IFNγ) gene signature generated from melanoma patient tumors that correlated with enhanced response to pembrolizumab across multiple cancer types [9]. While many clinical trials involving single-agent immunotherapy or combination therapies are being performed in NSCLC, a mechanistic understanding of determinants of response to these agents is still incomplete. These studies require preclinical models that recapitulate features of human lung cancer.

Our laboratory has employed an orthotopic immunocompetent mouse model to study how K-Ras mutant lung cancers respond to the immune system [12–15]. In this model lung cancer cells derived from C57BL/6J mice are implanted directly into the lungs of syngeneic mice. These cells form a primary tumor after 2-4 weeks that metastasizes to the other lung lobes, liver, brain, and mediastinum [16]. This model has the advantage that tumors develop in the appropriate tumor microenvironment (TME). In addition, the non-synonymous mutational burden in these tumors is comparable to human lung tumors, and significantly higher than genetically engineered mouse models [17], allowing for recognition by the adaptive immune system. We have previously demonstrated differential sensitivity of K-Ras mutant tumors to anti-PD-1/anti-PD-L1 therapy, with CMT167 tumors showing a strong inhibition and Lewis Lung Carcinoma (LLC) tumors being generally unresponsive [13]. The responsiveness of these tumors was also dependent on specific features of the lung TME. CMT167 tumors implanted subcutaneously were resistant to anti-PD-1 therapy whereas tumors in the lung were eliminated. Thus, this model allows us to define specific features of cancer cells that determine the response to immunotherapy. In this study we have focused on how cancer cell-intrinsic response to IFNγ affects the TME and response to anti-PD-1 therapy.

IFNγ is made predominantly by Natural Killer (NK), type 1 Innate Lymphoid (ILC1) and T cells [18, 19]. Since the 1990s it has been shown that IFNγ increases the immunogenicity of some tumors [20]. IFNγ binds to cell surface receptors (IFNGR1/ IFNGR2) on cancer cells resulting in activation of JAK1 and JAK2 and phosphorylation of STAT1 [20]. Activated STAT1 dimers translocate to the nucleus to initiate waves of transcription that can lead to enhanced MHC Class I and II presentation on tumor cells, and increased chemokine expression. Global loss of IFNγ is detrimental to tumor surveillance in mice, as *Ifnγ*^-/-^ mice develop tumors more quickly than their *Ifnγ*^+/+^ counterparts in the setting of carcinogen-induced or spontaneously arising tumors [21, 22]. Tumors that are insensitive to IFNγ can grow equally well in *Ifnγ*^-/-^ or *Ifnγ*^+/+^ mice, suggesting that host response does not completely alter the growth of these tumors [23]. Thus, it has been speculated that many tumors develop mutations in the IFNγ signaling pathway in order to evade the immune system. Recent studies have shown that approximately ~30% of both melanoma and lung carcinomas have at least one mutation in the IFNγ pathway including JAK1, IFNGR1, or IFNGR2 [20], and resistance to checkpoint inhibitors in patients is associated with JAK1/2 mutations [24].

We hypothesized that intrinsic differences in the responsiveness of cancer cells to IFNγ, distinct from other features of these cells, define the nature of the TME and control sensitivity of lung tumors to immunotherapy. In this study we demonstrated this by molecularly-altering responsiveness of murine lung cancer cells and defining changes in the TME that regulate responsiveness to anti-PD-1 therapy.

## Materials and Methods

### Cells

Murine Lewis Lung Carcinoma (LLC) cells expressing firefly luciferase were purchased from Caliper Life Sciences and maintained in DMEM (#10-017-CV, Corning) supplemented with 10% FBS, penicillin/streptomycin, and G418 (500ng/mL). LLC cells harbor a heterozygous K-Ras^G12V^ mutation [13]. CMT167 cells (gift of Dr. Alvin Malkinson, University of Colorado) were transduced with firefly luciferase and maintained in DMEM (#10-017-CV, Corning) with 10% FBS, penicillin/streptomycin, and G418 (500ng/mL) [16]. CMT167 cells harbor a K-Ras^G12C^ mutation [13]. Cell lines were confirmed mycoplasma-negative every two weeks and were last tested January 2019 (Lonza, # LT07-703). To maintain cellular phenotypes and to prevent cross-contamination of murine cell lines, cells were grown *in vitro* for less than 10 passages, and for only 2-3 weeks before use in *in vivo* experiments. Cell phenotypes were regularly assessed via proliferation assays, and EMT status. No phenotypic changes were observed during the course of these studies.

### Mice and Tumor Models

Wild-type C57BL/6J and green-fluorescent protein (GFP)-expressing mice [C57BL/6J-132Tg(UBC-GFP)30Scha/J] were obtained from Jackson Laboratory (Bar Harbor, ME). Dr. Haidong Dong (Mayo Clinic, Rochester, MN) provided PD-L1 knockout (KO) mice on a C57BL/6 background. Experiments were performed on 8-16 week old male and female mice. All mice were bred and maintained in the Center for Comparative Medicine at the University of Colorado Anschutz Medical Campus in accordance with established IACUC, U.S. Department of Health and Human Services Guide for the Care and Use of Laboratory Animals, and the University of Colorado Anschutz Medical Campus guidelines. For orthotopic lung tumors, an incision was made on the left lateral axillary line at the xyphoid process level, followed by removal of subcutaneous fat [25]. Tumor cells were suspended in 1.35 mg/mL Matrigel and Hank’s Buffered Saline Solution (1×10^5^ cells-LLC tumors; 5×10^5^ cells-CMT167 tumors in 40μl/injection) and injected into the left lung lobe through the rib cage with a 30-gauge needle [16]. For subcutaneous tumor cell implantation, animals were implanted with 1×10^6^ cells in the flank.

### Lentiviral Transduction and stimulation with IFNγ

Murine shRNA constructs were obtained from Sigma via the University of Colorado Functional Genomics Shared Resource (TRC1): Non-targeting control (SHC001V); shRNAs targeting *Socs1:* LLC-sh20 (TRCN0000067420), LLC-sh21 (TRCN0000067421); shRNAs targeting *Ifngr1*: CMT-sh68 (TRCN0000067368), and CMT-sh69 (TRCN0000067369). LLC or CMT167 cells were transduced with lentiviral particles generated from HEK293T cells transfected with shRNA vectors and lentiviral helper plasmids. Viral supernatant was collected at both 24 and 48 hours after transfection. Before transduction, LLC and CMT167 cells were pretreated with polybrene for 1 hour. During this time, polybrene was also added to viral supernatant generated from HEK293T cells and was filtered through a 0.45µm filter before media was placed on LLC or CMT167 cells. Stable cells were then selected by resistance to puromycin treatment (2 μg/mL). Pools of transduced cells were screened for degree of knockdown by mRNA and protein relative to parental cell lines and to the non-targeting control cells. For CMT167 transduced cells, knockdowns were subcloned and are subsequently represented as “CMT-sh68sc3” or “CMT-sh69sc2”. For *in vitro* experiments, cells were treated with recombinant murine IFNγ (0-100 ng/mL) (PeproTech #315-05) followed by isolation of protein and/or RNA for immunoblotting and quantitative real-time-PCR (qRT-PCR).

### Immunoblotting

Cells were washed 3X with PBS, followed by lysis with MAPK Buffer(50 mM β-glycerophosphate, pH 7.2, 0.5% Triton X-100, 5 mM EGTA, 100 μM sodium orthovanadate, 1 mM dithiothreitol, and 2 mM MgCl2) and a protease inhibitor cocktail from Sigma (Sigma #P8340). 10-40 μg of total protein was fractionated by SDS-polyacrylamide gel electrophoresis and transferred to PVDF membranes. Antibodies used were: pSTAT1 (Y701) Cell Signaling #9167S (1:500-1:1000); STAT1, Cell Signaling #9172S (1:1,000-1:1,500); SOCS1, Abcam #ab3691 (1:300-1:500); IFNGR1 (Interferon Gamma Receptor Alpha), Lifespan Biosciences #LS-C33-4260 (1:300-1:500); IFNGR2 (Interferon Gamma Receptor Beta/AF-1), Abcam #77246 (1:300-1:500); β-ACTIN, Sigma #A5441 (1: 5,000-1:10,000); Rabbit HRP, Jackson Immuno Research #111-035-144 (1: 5,000-1:10,000); Mouse HRP, Santa Cruz #2c-2005 (1: 5,000-1:10,000).

### Quantitative Real-time-PCR

Total RNA from cultured cells was isolated using the RNeasy Mini Kit (Qiagen), followed by reverse transcription with 1μg of total RNA/sample (qScript cDNA Synthesis Kit, QuantaBio). qRT-PCR was conducted on the myIQ Real Time PCR Detection System (BioRad) using Power SYBR Green PCR Master Mix (Applied Biosystems). Relative message levels of each gene were normalized to the housekeeping gene, *Actb* (shown as Absolute Values, or SQ Values). Primers: Murine *Socs1*, F:5’-CTGCGGCTTCTATTGGGGAC-3’, R:3’-AAAAGGCAGTCGAAGGTCTCG-5’; Murine *Cxcl9*, F:5’-GAGCAGTGTGGAGTTCGAGG-3’, R:3’-TCCGGATCTAGGCAGGTTTG-5’; Murine *Ciita*, F:5’-TGCGTGTGATGGATGTCCAG-3’, R: 3’-CCAAAGGGGATAGTGGGTGTC-5’; Murine *Cxcl10*, F:5’-GGATGGCTGTCCTAGCTCTG-3’, R:3’-TGAGCTAGGGAGGACAAGGA-5’; Murine *Ifngr1*, F: 5’-TACAGGTAAAGGTGTATTCGGGT-3’, R:3’-ACCGTGCATAGTCAGATTCTTTT-5’; Murine *Cd274* (PD-L1),F:5’-TGCTGCATAATCAGCTACGG-3’, R:3’-GCTGGTCACATTGAGAAGCA-5’; Murine *Actb*, F: 5’-GGCTGTATTCCCCTCCATCG-3’, R: 3’-CCAGTTGGTAACAATGCCATG-5’.

### Cxcl9 ELISA

Tumor cells were treated for 48 hours with 100 ng/mL IFNγ *in vitro*. Media was collected, spun down to remove floating cells, and ELISA was performed on supernatant according to manufacturer’s protocol. 50µL of conditioned media was used per replicate. ELISA: R&D Systems #DY492.

### PD-L1 Expression of Cancer Cells by Flow Cytometry

Tumor cells were treated for 18 hours with 100 ng/mL IFNγ and/or 1μM Ruxolitinib (LC Laboratories #R-6688, 1µM) *in vitro*. Cells were scraped, washed with PBS, and re-suspended in antibody solution. Flow cytometry was performed on the Yeti instrument and analyzed using Kaluza software as part of the University of Colorado Cancer Center Flow Cytometry Core. Antibodies: Anti-Mouse PD-L1-PE, eBioscience #12-5982-81 (1:200); Ghost 510 Viability Dye, Tonbo Biosciences #13-0870-T100 (1:200).

### Anti-PD-1 Treatment

Tumor-bearing mice were intraperitoneally injected twice weekly with either an IgG2a isotype control antibody, or an anti-PD-1 antibody (BioXCell) at a dose of 200 Antibodies: Anti-Mouse IgG2a, BioXCell #BE0089 Clone 2A3; Anti-Mouse PD-1, BioXCell #BE0146 Clone RMP1-14.

### Immunofluorescence

Tumor-bearing lungs were perfused with 20 U/mL of PBS/Heparin followed by inflation, fixed overnight in 10% formalin and maintained in 70% ethanol until paraffin embedding. 4μm thick sections cut from FFPE tissue blocks were deparaffinized, rehydrated and stained with 0.1% Sudan Black B (Sigma) in 70% ethanol. Slides were heated in a citrate antigen retrieval solution for 2 hours at 100°C and quenched with 10 mg/mL Sodium Borohydride. Slides were blocked with a mixture of goat serum, Superblock (SkyTek Laboratories) and 5% BSA overnight. Slides were incubated with primary antibodies in a 1:1 mixture of 5% BSA and SuperBlock for 1 hour, followed by incubation with secondary antibodies for 40 minutes. Slides were coverslipped with Vectashield with Dapi. Hematoxylin and eosin (H&E) stains were performed on one section per tumor by the University of Colorado Denver’s Histology Shared Resource Core. For quantitation of T cells, at least three non-serial tumor sections per animal (6 animals per experimental condition) were examined. The mean number of CD3^+^/CD4^+^/CD8^+^ T cells was obtained from the average of 6 random 40X tumor fields per section using two blinded observers. Antibodies/Reagents used: Anti-Mouse CD3e, ThermoScientific #MA5-14524 Clone SP7 (1:100); Anti-Mouse CD4, eBioscience #14-9766-82 (1:50); Anti-Mouse CD8, eBioscience #14-0808-82 (1:100); AF594 Goat anti-rabbit IgG, Lifetech #A11037(1:1,000); AF488 Goat anti-rat, Lifetech #A11006 (1:1,000); Vectashield with Dapi, Vector #H-1200. Microscope: Nikon Eclipse Ti-S #TI-FLC-E at 40X/0.75, ∞/0.17 WD 0.72. Camera: Zyla scMOS, Andor #DG-152VC1E-FI. Acquisition Software: NIS Elements 64-Bit AR 4.60.00. Data Analysis: FIJI.

### In-situ Hybridization

Sections (4μm) of lung tumor tissue underwent deparaffinization, followed by treatment with RNAscope hydrogen peroxide for 10 minutes at RT, and 1X Target Retrieval Reagent at 99ºC for 15-30 minutes. Slides were then treated with RNAscope Protease Plus for 15-30 minutes at 40ºC in the HybEZ Oven. Following pretreatment, slides were treated using the RNAscope 2.5 HD Detection Reagent-BROWN kit per the manufacturer’s protocol. The signal was detected for either the negative control probe (dapB), or murine *Cxcl9*. Reagents: RNAScope Target Retrieval Reagents, Advanced Cell Diagnostics (ACD) #322000; RNAScope Wash Buffer Reagents, ACD #310091; RNAScope 2.5 HD Detection Reagent-BROWN, ACD #322310; RNAScope H2O2 & Protease Plus Reagents, ACD #322330; RNAScope Negative Control Probe_dapB, ACD #310043; RNAScope Probe Mm-Cxcl9, ACD #489341; HybEZ II Oven, ACD #321710/321720; Humidity Control Tray, ACD #310012; EZ-Batch Wash Tray, ACD #310019; EZ-Batch Slide Holder, ACD #310017. Microscope/Camera: Olympus BX41 System at 40X/0.65, ∞0.17/FN22. Acquisition Software: SPOT. Data Analysis: FIJI.

### Flow Cytometry

Mice were sacrificed between 2-4 weeks post-tumor cell injection. Tumor-bearing left lung lobes were excised, mechanically dissociated, and incubated at 37°C for 30 minutes with collagenase type II (8480 U/ml, Worthington Biochemical), elastase (7.5mg/ml, Worthington Biochemical), and soybean trypsin inhibitor (2mg/ml, Worthington Biochemical). After which, single-cell suspensions were made and filtered through 70 µm cell strainers (BD), subjected to red blood cells lysis using hypotonic buffer (0.15 mM NH4Cl, 10 mM KHCO3, 0.1 mM Na2EDTA, pH 7.2), and filtered again through 40 µm cell strainers (BD)[12]. For the “T Cell Phenotypic Panel”, single cell suspensions were stained for 30 minutes at room temperature, followed by fixation and permeabilization overnight, and intracellular stains for 2 hours at 4°C the following day. For the “T Cell Stimulation Panel”, single cell suspensions were stimulated with Brefeldin A solution, Monensin solution, and a cell stimulation cocktail (PMA/Ionomycin) for 5 hours at 37°C. Afterwards, single cell suspensions were stained with cell surface stains, fixed and permeabilized overnight, and finally stained with intracellular stains the following morning (as the T Cell Phenotypic Panel). Samples were run at the University of Colorado Cancer Center Flow Cytometry Core using the Gallios system (Beckman Coulter). The gating strategy involved excluding debris and cell doublets by light scatter, as well as dead cells by a cell viability dye. All data was analyzed using Kaluza Software (Beckman Coulter). Antibodies and Reagents: Foxp3 Staining Buffer Set, eBioscience #00-5523-00; Brefeldin A, Biolegend #420601; Monensin, Biolegend #420701; Cell Stimulation Cocktail, eBioscience #00-4970-93; Anti-Mouse PD-1-PE, eBioscience #12-9981-81 (1:200); Anti-Mouse CD69-PECy7, eBioscience #25-0691-81 (1:200); Anti-Mouse CD45-AF700, eBioscience #56-0451-82 (1:50); Anti-Mouse IA/IE Dazzle 594, Biolegend #107648 (1:250); Anti-Mouse CD3e-PerCP Cy5.5, eBioscience #45-0031-82 (1:200); Anti-Mouse CD4-EF450eBioscience #48-0042-82 (1:200); Anti-Mouse CD8a-APC EF780, eBioscience #47-0081-82 (1:200); Murine Fc Block, eBioscience #14-0161-86 (1:100); V500 rat anti-mouse CD4, BD #560782; Aqua Viability Dye, ThermoFisher #L34957 (1:200); Rat Anti-Mouse Isotype for IFNγ-AF488, eBioscience #53-4301-80 (1:80); Anti-Mouse IFNγ-AF488, eBioscience #53-7311-82(1:80); Rat Anti-Mouse Isotype for PerCP Cy5.5-TNFα, BD #560537(1:80); Anti-Mouse PerCP Cy5.5-TNFα, BD #560659(1:80); VersaComp Antibody Capture Bead Kit, Beckman Coulter #B22804. Each replicate consisted of 3 tumor bearing lung lobes pooled for a total of 9 mice per experimental condition.

### CyTOF Analysis

Single cell suspensions prepared as above were treated with benzonase nuclease (Sigma #E1014, 1:10,000), stained with cisplatin, and fixed for sample barcoding (Fluidigm). Samples were then combined into one tube, followed by incubation with an Fc receptor blocking antibody, primary surface antibodies, and secondary surface staining. Cells were then fixed and permeabilized overnight, followed by intracellular stains the next day. After staining, cells were suspended in Intercalator [26]. Single cell suspensions were run on the Helios Mass Cytometer as part of the University of Colorado Cancer Center Flow Cytometry Core. Antibodies and Reagents: 89Y-CD45, Fluidigm, Clone 30-F11; 141Pr-Gr1 (Ly6C/Ly6G), Fluidigm, Clone RB6-8C5; 142Nd-CD11c, Fluidigm, Clone N418; 143Nd-GITR, Fluidigm, Clone DTA1; 144Nd-MHC class I, Fluidigm, Clone 28-14-8; 145Nd-SiglecF-PE/anti-PE, BD, Clone E50-2440/ Fluidigm, Clone PE001; 146Nd-CD8a, Fluidigm, Clone 53-6.7; 147Nd-p-H2AX[Ser139], Fluidigm, Clone JBW301; 148Nd-CD11b, Fluidigm, Clone M1/70; 149Sm-CD19, Fluidigm, Clone 6D5; 150Nd-CD25, Fluidigm, Clone3C7; 151Eu-CD64, Fluidigm, Clone X54-5/7.1; 152Sm-CD3e, Fluidigm, Clone 145-2C11; 153Eu-PD-L1, Fluidigm, Clone 10F.9G2; 154Sm-CTLA4, Fluidigm, Clone UC10-4B9; 155Gd-IRF4, Fluidigm, Clone 3E4; 156Gd-CD90.2(Thy-1.2), Fluidigm, Clone 30-H12; 158Gd-FoxP3, Fluidigm, Clone FJK-16s; 159Tb-PD-1, Fluidigm, Clone RMP1-30; 160Gd-CD80/86-FITC/anti-FITC, BD, Clone 16-10A1/BD Clone BL1/Fluidigm, Clone FIT22; 161Dy-INOS, Fluidigm, Clone 4B10; 162Dy-Tim3, Fluidigm, Clone RMT3-23; 163Dy-CXCR3-APC/anti-APCCXCR3-173, Biologend/Fluidigm, Clone APC003; 164Dy-IkBa, Fluidigm, Clone L35A5;165Ho-Beta-catenin (active), Fluidigm, Clone D13A1; 166Er-Arg1, Fluidigm, Clone 6D5; 167Er-NKp46, 2 Fluidigm, Clone 9A1.4; 168Er-Ki-67; Fluidigm, Clone Ki-67; 169Tm-Ly-6A/E (Sca-1), Fluidigm, Clone D7; 170Er-CD103-Biotin/anti-Biotin, Biolegend, Clone 2E7/Fluidigm, Clone 1D4-C5; 171Yb-CD44, Fluidigm, Clone IM7; 172Yb-CD4, Fluidigm, Clone RM4-5; 173Yb-CD117 (ckit), Fluidigm, Clone 2B8; 174Yb-Lag3, Fluidigm, Clone M5/114.15.2;175Lu-CD127, Fluidigm, Clone A7R34; 176Yb-ICOS, Fluidigm, Clone 7E.17G9; 191Ir, 193Ir-Intercalator, Cell-ID; 195Pt Cisplatin 5 uM, Cell-ID; 140Ce, 151Eu, 153 Eu, 165Ho, 175Lu Normalization Beads; Cell ID 20-Plex Pd Barcoding Kit, Fluidigm,#201060 (102, 104, 105, 106, 108, 110 Pd Bar Codes); Benzonase, Sigma #E1014-5KU(1:10,000) in HBSS. Each replicate consisted of 3 tumor bearing lung lobes (or naïve lungs) pooled for a total of 9 mice per experimental condition.

### PhenoGraph Analysis Methods

Software for data analysis included R studio (Version 1.0.136), downloaded from the official R Web site (https://www.r-project.org/); the cytofkit package (Release 3.6), downloaded from Bioconductor (https://www.bioconductor.org/packages/release/bioc/html/cytofkit.html); Excel 15.13. 14, FlowJo 10.2, GraphPad Prism 7, and Adobe Illustrator CC 2017. Samples were normalized using NormalizerR2013b_MacOSX, downloaded from the Nolan laboratory GitHub page (https://github.com/nolanlab). The normalized files were then debarcoded using SingleCellDebarcoderR2013b_MacOSX, downloaded from the Nolan laboratory GitHub page (https://github.com/nolanlab). Debarcoded and normalized data were subjected to traditional Boolean gating in FlowJo, identifying viable singlet events (191Ir+, 193Ir+, 195Pt-). These events were exported for downstream analysis. All viable singlet (19Ir +, 193Ir +, 195Pt +) events were imported into cytofkit analysis pipeline and 39 markers were selected for clustering. The merge method ‘min’ was selected (12,255 events from each file used for clustering) and the files were transformed via the cytofAsinh method. Then files were clustered with the PhenoGraph algorithm and tsne was selected as the visualization method. PhenoGraph identified 35 unique clusters. These results were visualized via the R package “Shiny” where labels, dot size, and cluster color were customized according to cluster identity or phenotype. Plots were examined for expression of various cellular markers (parameters). The algorithm produced multiple .csv files, the files “cluster median data” and “cluster cell percentage” which were utilized to determine cluster frequency and phenotype.

### RNA-Sequencing Analysis of Cancer Cells Recovered from GFP-transgenic Mice

GFP-expressing transgenic mice were implanted with 10^5^ cells as described above. After 2-3 weeks of tumor growth, single cell suspensions of tumor-bearing lung lobes were prepared containing a mixture of GFP-negative cancer cells and GFP-positive host cells. GFP-negative cancer cells were sorted using the MoFlo XDP cell sorter with a 100μm nozzle (Beckman Coulter) as part of the University of Colorado Cancer Center Flow Cytometry Shared Resource. The sorting strategy excluded dead cells (via DAPI staining) and cell doublets by light scatter. Total RNA was isolated via the RNeasy Plus Mini Kit (Qiagen). CMT167, LLC, LLC-NT, and LLC-sh21 cells were recovered from 3-5 pools of mice consisting of at least four GFP-expressing mice per single pool. Total RNA was also isolated from cells in culture at the time of injection. Preparation of the RNA-Sequencing (RNA-Seq) library was done at the University of Colorado Cancer Center Genomics and Microarray Shared Resource. RNA libraries were constructed using an Illumina TruSEQ stranded mRNA Sample Prep Kit and sequencing was performed using an Illumina HiSEQ 4000 System. Reads from RNA-Sequencing were processed and aligned to a mouse reference genome (UCSC Mus musculus reference genome build mm10) via the TopHat v2 software [15]. The aligned read files were then processed by Cufflinks v2.0.2 software in order to determine the relative abundance of mRNA transcripts[15]. Reads are portrayed as Fragments Per Kilobase of exon per million fragments mapped (FPKM). For pathway analysis of final FPKM files, various analysis platforms including KEGG and DAVID were used to determine the most highly enriched pathways between experimental conditions.

### Statistical Analysis

Statistical Analyses were performed using the GraphPad Prism 7/8 software. Data are presented as mean α SEM. A *one-or two-way ANOVA* was used to compare differences in more than two groups. A *student’s t-test* was used to compare differences between two groups in data with a normal distribution. In all circumstances, *p-values ≤0.05* were considered significant (*p<0.05, **p<0.01, ***p<0.001, ****p<0.0001).

## Results

### LLC Cells Exhibit a Blunted Response to IFNγ In Vitro and In Vivo Compared to CMT167

We hypothesized that the differential response of CMT167 versus LLC orthotopic tumors to anti-PD-1 therapy was mediated at least in part through inherent differences in the cancer cells and how they respond to signals coming from the tumor microenvironment (TME). To define these changes, we recovered cancer cells from orthotopically implanted tumors, and compared their transcriptional profile to identical cancer cells grown *in vitro*. CMT167 or LLC cells were injected into the lungs of transgenic GFP-expressing C57BL/6J mice. After tumors were established, the GFP-negative cancer cell population was recovered by fluorescence activated cell sorting (FACS) of single-cell suspensions made from tumor-bearing lungs. RNA isolated from recovered cells and from identical cells grown *in vitro* was analyzed by RNA-Seq. We used pathway analysis to identify pathways that were differentially induced *in vivo* in the two cell lines. From this analysis, we determined that the interferon-gamma (IFNγ) signaling pathway was selectively upregulated in CMT167 tumors compared to LLC tumors (**Figure 1A**), suggesting that these cells have a differential response to IFNγ. Examination of RNA-Seq data revealed no detectable mutations in the IFN signaling pathways in either cell line (data not shown). Both cells expressed comparable levels of IFNγ receptors, although expression was higher in CMT167 cells (**Supplemental Figure 1A-C**) and JAK/STAT machinery, implicating other potential alterations in intracellular signaling. To validate our RNA-Seq data, we compared the responsiveness of these two cell lines to IFNγ treatment *in vitro*. Stimulation with recombinant murine IFNγ showed a significantly greater induction of 4 IFNγ response genes (*Cxcl9, Cxcl10*, *Cd274* and *Ciita*) in CMT167 cells compared to LLC cells (**Figure 1B-E**). Additionally, CMT167 cells showed a more robust and sustained induction of phospho-STAT1 (p-STAT1), and total STAT1 levels upon treatment with IFNγ compared to LLC cells (**Figure 1F**). Collectively these data suggest that responsiveness of the cancer cells to IFNγ signaling is associated with sensitivity to anti-PD-1 therapy in our model.

**Figure 1.**
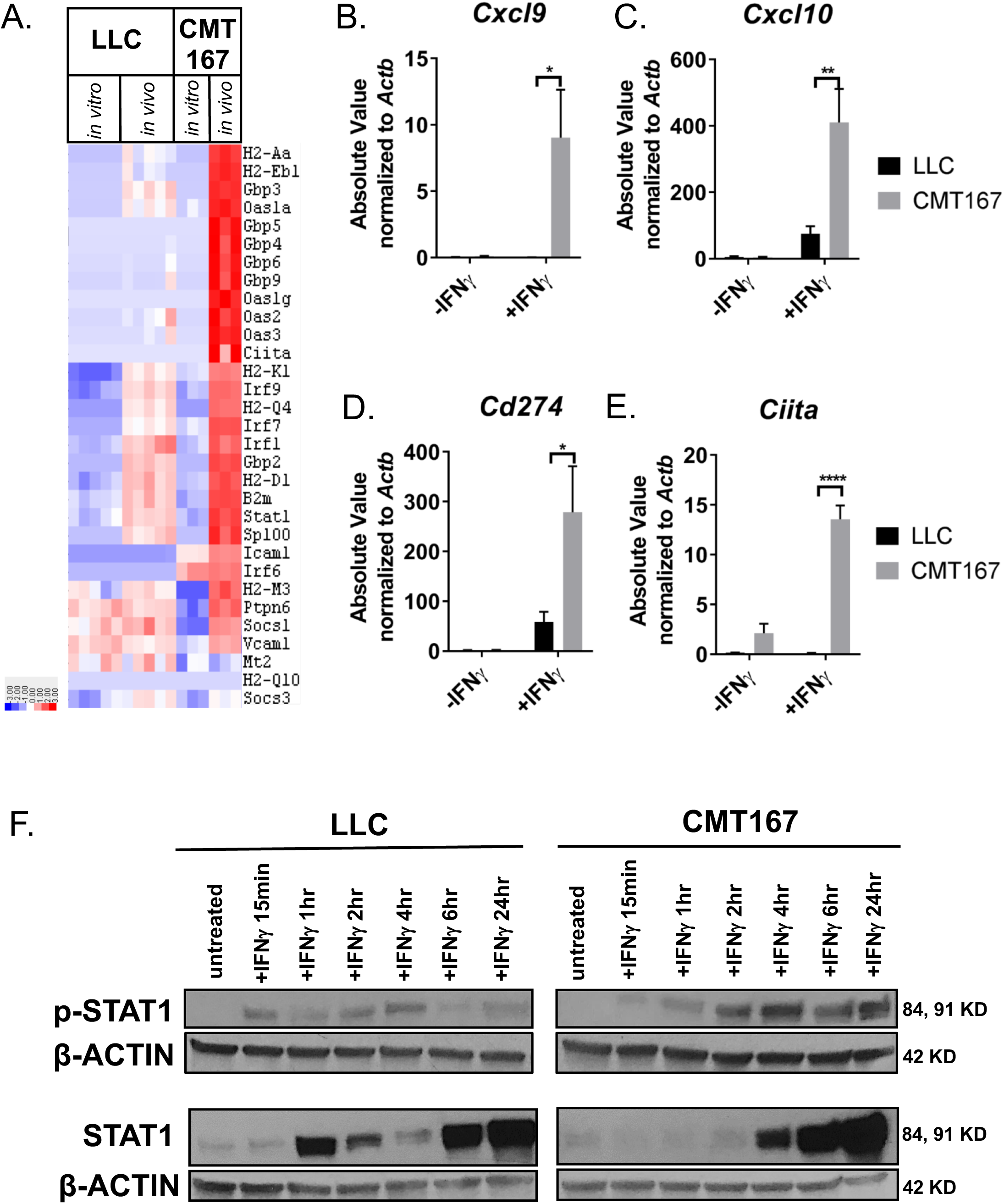
LLC Cells Exhibit a Blunted Response to IFNγ In Vitro and In Vivo Compared to CMT167. CMT167 or LLC cells were injected into the left lung lobe of transgenic GFP-expressing C57BL/6J mice and grown for either 2 (LLC) or 3 weeks (CMT167). Single cell suspensions of the tumor-bearing lobe were flow sorted to recover GFP-positive (host cells) and GFP-negative (tumor cells). RNA was isolated from the GFP-negative cells (*in vivo* condition) and from identical cells grown in passage (*in vitro condition)* and analyzed by RNA-seq. CMT167 samples had 3 experimental replicates per *in vitro* and *in vivo* conditions, with five tumor bearing lung lobes pooled per *in vivo* experimental replicate (15 mice used total). LLC samples had 5 experimental replicates per *in vitro* and *in vivo* conditions with four tumor bearing lung lobes pooled per *in vivo* experimental replicate. (A) KEGG Pathway Analysis of RNA-Seq data from the *in vitro* and *in vivo* experimental conditions showing that the CMT167 line expresses a robust IFNγ signature *in vivo*, while the LLC line has a much more blunted response. LLC or CMT167 cells were treated with ±10ng/mL of IFNγ *in vitro* for 24 hours followed by isolation of RNA and qRT-PCR. mRNA levels of (B) *Cxcl9*, (C) *Cxcl10*, (D) *Cd274*, and (E) *Ciita* are shown as Absolute Values (SQ Values) normalized to the housekeeping gene *Actb*. Statistics compare IFNγ treated LLC and CMT167 cells. (F) Immunoblots of LLC or CMT167 cells treated with ±10ng/mL of IFNγ *in vitro* for a time course ranging from 15 minutes to 24 hours, showing p-STAT1 and total STAT1 expression compared to the housekeeping gene β-ACTIN. Error bars represent the mean of the data ±SEM after a two-way ANOVA (1B-E) (*p<0.05, **p<0.01, ***p<0.001, ****p<0.0001).

### Silencing Ifngr1 in CMT167 Confers Decreased Response to IFNγ and Resistance to Anti-PD-1 Therapy

To confirm that the responsiveness to IFNγ of CMT167 cells affects response to checkpoint inhibitors, *Ifngr1* was silenced in CMT167 cells using 2 separate shRNAs against murine *Ifngr1* and a non-targeting control vector. Expression of *Ifngr1* was decreased by approximately 80% with both shRNA constructs (CMT-sh68sc3, CMT-sh69sc2) compared to non-targeting control (CMT-NT) (**Figure 2A**). Importantly, induction of downstream IFN response genes after IFNγ treatment (*Cxcl9, Cxcl10, Cd274, and Ciita*,) was markedly inhibited in both shRNA knockdowns (**Figure 2B-E**). This was also associated with decreased induction of p-STAT1 in response to IFNγ stimulation (**Figure 2F**). We selected one knockdown, CMT-sh68sc3, for *in vivo* studies. Equal numbers of CMT-sh68sc3 or CMT-NT were implanted into the lungs of syngeneic WT mice that were treated with either a control IgG2a antibody or an antibody targeting PD-1, starting 7 days post tumor cell injection. After 4 weeks, we found that CMT-NT tumors treated with anti-PD-1 were almost completely eliminated similar to the published CMT167 parental line (**Figure 2G**) [13]. However, treatment of CMT-sh68sc3 *Ifngr1* KD tumors with anti-PD-1 had no significant effect on tumor size (**Figure 2G**). We previously reported anti-PD-1 treatment of CMT167 tumors results in nests of infiltrating T cells associated with tumor elimination [13]. While we observed similar patterns of T cell infiltration in CMT-NT tumors, this was not observed in the CMT-sh68sc3 tumors (**Supplemental Figure 1D-E**). These data indicate that the IFNγ responsiveness of CMT167 cells is critical for their response to immunotherapy.

**Figure 2.**
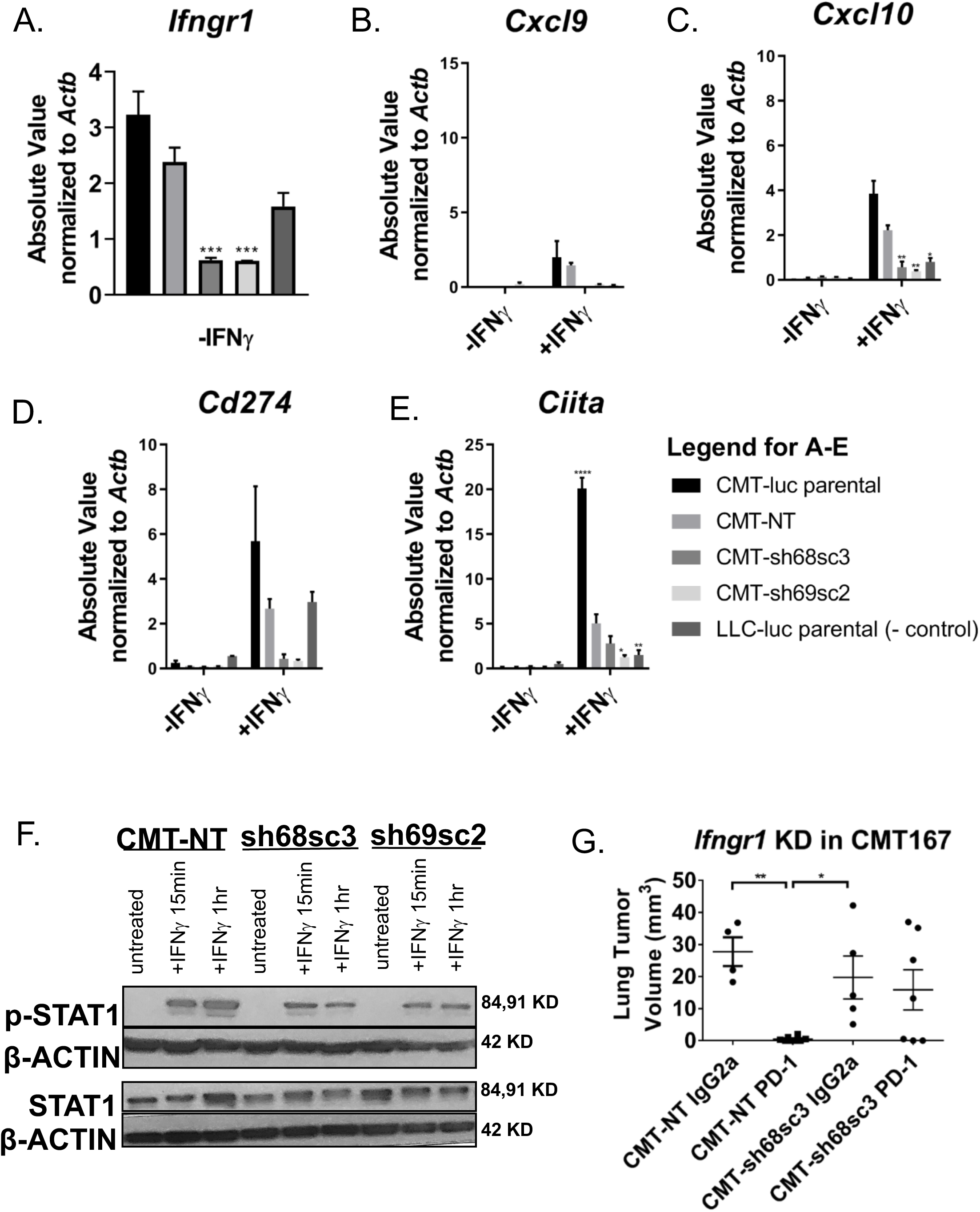
Silencing Ifngr1 in the CMT167 Line Confers Decreased Response to IFNγ In Vitro and In Vivo. Two separate shRNAs targeting *Ifngr1* (CMT-sh68, CMT-sh69) and a non-targeting control vector (CMT-NT) were transduced into CMT167 cells expressing luciferase. CMT167 cells were then screened for functional and stable knockdown of *Ifngr1* after 10 days of puromycin selection and subcloning of shRNA pools (CMT-sh68sc3, CMT-sh69sc2). Cells were treated with ± 10ng/mL IFNγ for 24 hours followed by isolation of RNA and qRT-PCR. mRNA levels of (A) *Ifngr1*, (B) *Cxcl9*, (C) *Cxcl10*, (D)*Cd274* and (E) *Ciita* are shown as Absolute Values (SQ Values) normalized to the housekeeping gene *Actb*. Statistics compare the CMT-NT cell line to other cell lines with or without treatment. (F) Immunoblots showing p-STAT1, total STAT1, and β-ACTIN levels of the CMT-NT, CMT-sh68sc3, and CMT-sh69sc2 cell lines ±IFNγ after 15 minutes or 1 hour *in vitro*. (G) CMT-NT or CMT-sh68sc3 (*Ifngr1* knockdown) cells were orthotopically injected into the lungs of syngeneic mice, established for 7 days, then were treated with either an isotype control antibody (IgG2a) or an anti-PD-1 antibody for 3 weeks followed by terminal sacrifice at 4 weeks post tumor cell injection. Primary tumor volume was assessed using digital calipers. Error bars represent the mean of the data ±SEM after a one-way ANOVA (2A,2G), or a two-way ANOVA (2B-E) (*p<0.05, **p<0.01, ***p<0.001, ****p<0.0001).

### Silencing Socs1 in the LLC Line Confers Increased Response to IFNγ *In Vitro*

Since the lack of an IFNγ response in LLC cells is not due to lack of receptor expression, we examined differences in expression of putative regulators of the IFNγ pathway between the responsive CMT167 and unresponsive LLC cell lines. We determined that at baseline, LLC cells expressed markedly higher levels of *Socs1*, or suppressor of cytokine signaling 1, which is a critical negative regulator of interferon signaling (**Figure 3A**) [27, 28]. We confirmed that LLC cells expressed higher levels of both SOCS1 protein and mRNA relative to CMT167 cells *in vitro* (**Figure 3B-C**). These data led us to hypothesize that high baseline levels of *Socs1* mediate the unresponsiveness of LLC cells to IFNγ. Thus, silencing *Socs1* should increase LLC cells’ response to IFNγ *in vitro* and potentially alter their response to checkpoint inhibitors *in vivo*.

**Figure 3.**
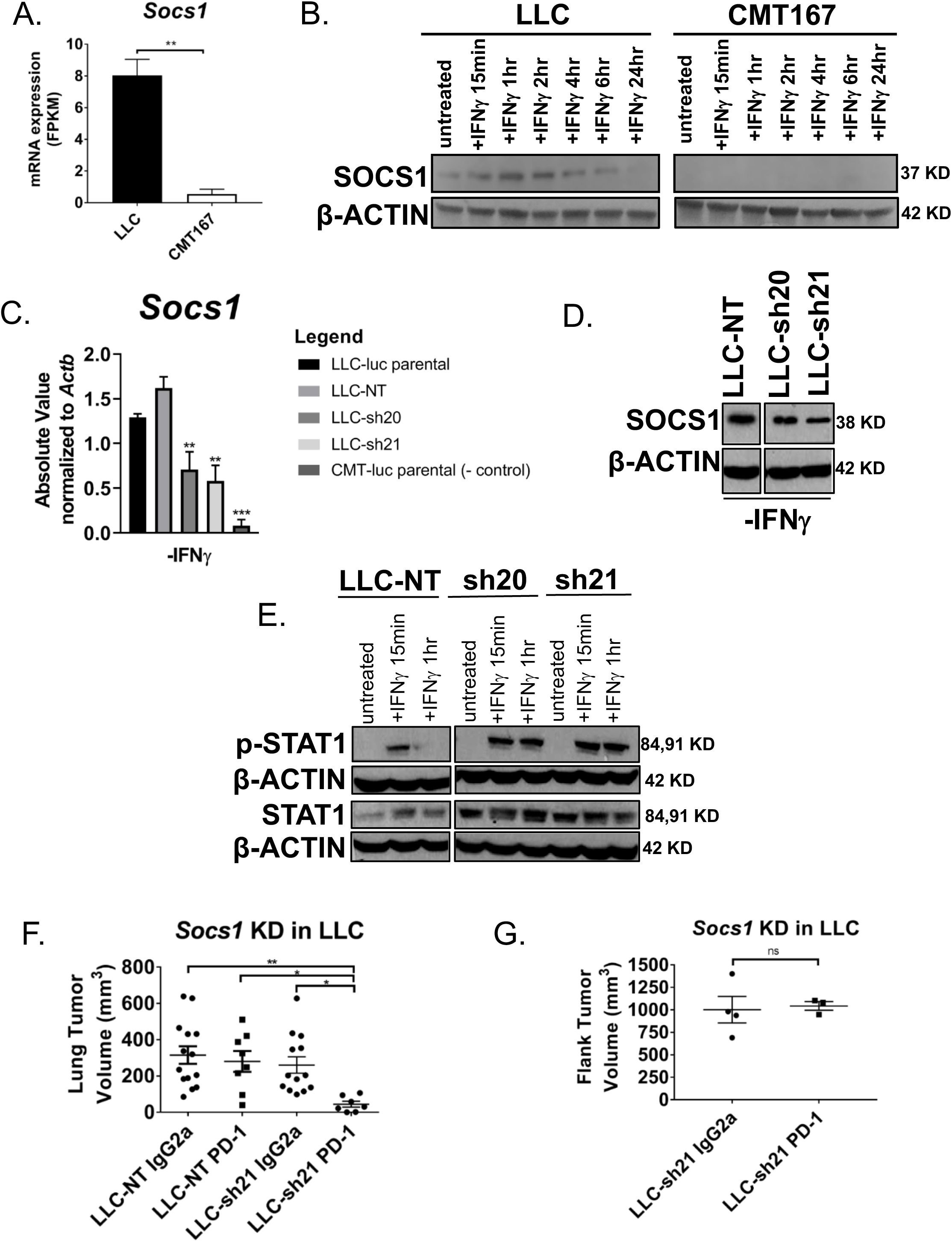
Silencing Socs1 in the LLC Line Confers Increased Response to IFNγ In Vitro and In Vivo. (A) *In vitro* mRNA expression of *Socs1* in FPKM as assessed by RNA-Seq in LLC and CMT167 cells. (B) Immunoblots of LLC or CMT167 cells treated with ±10ng/mL of IFNγ *in vitro* for a time course ranging from 15 minutes to 24 hours, showing SOCS1 expression relative to the housekeeping gene β-ACTIN. (C) mRNA expression of *Socs1* via qRT-PCR shown as Absolute Values (SQ Values) normalized to the housekeeping gene *Actb* for LLC cells transduced with shRNA constructs(LLC-sh20, LLC-sh21), an non-targeting shRNA (LLC-NT), and compared to parental LLC-luc or CMT167-luc cells. Statistics compare the LLC-NT line to other cell lines. (D) Immunoblot for SOCS1 relative to β-ACTIN protein levels in the LLC-NT, LLC-sh20, and LLC-sh21 cell lines *in vitro*. (E) Immunoblots showing p-STAT1, total STAT1, and β-ACTIN levels of the LLC-NT, LLC-sh20, and LLC-sh21 cell lines ±IFNγ at 15 minutes or 1 hour *in vitro.* (F) LLC-NT or LLC-sh21 (*Socs1* knockdown) cells were orthotopically injected into the lungs of syngeneic mice, established for 7 days, then were treated with either an isotype control antibody (IgG2a) or an anti-PD-1 antibody for 2 weeks followed by terminal sacrifice at 3 weeks post tumor cell injection. Primary tumor volume of lung tumors was assessed using digital calipers. (G) LLC-sh21 cells were injected into the flanks of syngeneic mice, established for 7 days, then were treated as in Figure F. Primary tumor volume of subcutaneous tumors was assessed with digital calipers. Error bars represent the mean of the data ±SEM after a student’s t-test (3A,3G), or a one-way ANOVA (3C, 3F) (*p<0.05, **p<0.01, ***p<0.001, ****p<0.0001).

*Socs1* expression in LLC cells was silenced using 2 separate shRNAs and a non-targeting control construct (LLC-sh20, LLC-sh21, LLC-NT). As anticipated, knockdown variants had decreases in *Socs1* mRNA and protein (**Figure 3C-D**). To assess the functionality of our knockdowns, we treated LLC-NT versus *Socs1*-KD cells with IFNγ. We found that induction of multiple IFNγ response genes (*Cxcl9, Cxcl10, Cd274, and Ciita)* was enhanced in both knockdowns compared to LLC-NT cells (**Supplemental Figure 2A-D**); there was no change in expression of *Ifngr1* levels (**Supplemental Figure 2E**). Compared to LLC-NT cells, both knockdowns exhibited enhanced STAT1 signaling at early and late time points as determined by expression of p-STAT1 (**Figure 3E**, **Supplemental Figure 2F**). We chose LLC-sh21 cells for further studies and validated that upon IFNγ stimulation, they had increased CXCL9 (measured by ELISA) and PD-L1 (measured by flow cytometry) *in vitro* relative to LLC-NT cells (**Supplemental Figure 2G-H**). Induction of PD-L1 protein by IFNγ was completely inhibited by Ruxolitinib, an inhibitor of JAK1/JAK2 in the LLC-sh21 cells, indicating that the induction of this gene was JAK/STAT dependent (**Supplemental Figure 2H**). These data collectively indicate that LLC cells are refractory to IFNγ signaling due to high basal levels of *Socs1*. Additionally, *Socs1* knockdown in LLC cells sensitizes them to IFNγ by increasing the magnitude and duration of JAK/STAT signaling.

### Socs1 KD Tumors Show Enhanced Response to Anti-PD-1 Therapy

To determine if altering the sensitivity of LLC cells to IFNγ affects tumor growth *in vivo* and responsiveness to anti-PD-1, we implanted LLC-NT or LLC-sh21 cells into the lungs of syngeneic WT mice. Tumors were allowed to establish for 1 week and were then treated with either an anti-PD-1 antibody or an isotype control antibody (IgG2a) for 2 weeks (as above). Similar to the parental LLC line as previously published [13], there was no significant difference in primary tumor volume between the LLC-NT tumors treated with anti-PD-1 or isotype control after 3 weeks (**Figure 3F**). However, in mice harboring LLC-sh21 tumors, treatment with the anti-PD-1 antibody decreased primary tumor volume by over 80%, a statistically significant difference compared to all the other experimental groups (**Figure 3F**). To determine if these effects were specific to the lung TME, we analyzed the response of LLC-sh21 cells implanted subcutaneously to anti-PD-1 therapy. Unlike what was observed in orthotopic lung tumors, subcutaneous LLC-sh21 tumors were resistant to anti-PD-1 therapy (**Figure 3G**). This is similar to our previous data showing that the sensitivity of CMT167 tumors to anti-PD-1 therapy was specific to tumors implanted into the lung, while identical cells implanted subcutaneously were resistant [13]. These data suggest enhanced responsiveness of LLC tumors to anti-PD-1 is dependent on critical features of the lung TME that are absent in subcutaneous models.

### Socs1 KD Tumors Show Alterations in Multiple Populations

To define changes in the TME that are dependent on the IFNγ-responsiveness of the LLC cells, we used mass cytometry (CyTOF) to characterize both CD45^+^ and CD45^-^ populations from LLC-NT and LLC-sh21 tumors prior to treatment with anti-PD-1 using a panel of 39 different antibodies. Three independent isolations using 3 mice/isolation were analyzed. There were no significant differences in tumor size between LLC-NT and LLC-sh21 at this time point (**Supplemental Figure 3A**). Using the Phenograph algorithm, which allows unbiased clustering of events based on cellular distribution and phenotype, we identified 35 distinct clusters in the naïve, LLC-NT and LLC-sh21 experimental conditions [29]. **Figure 4A** depicts the tSNE plot for all samples, with clusters colored by phenotype. Phenotypes were defined based on the expression level of cellular markers (parameters) (**Supplemental Figure 3B**). No significant differences were noted between the replicates (**Supplemental Figure 3C**).

**Figure 4.**
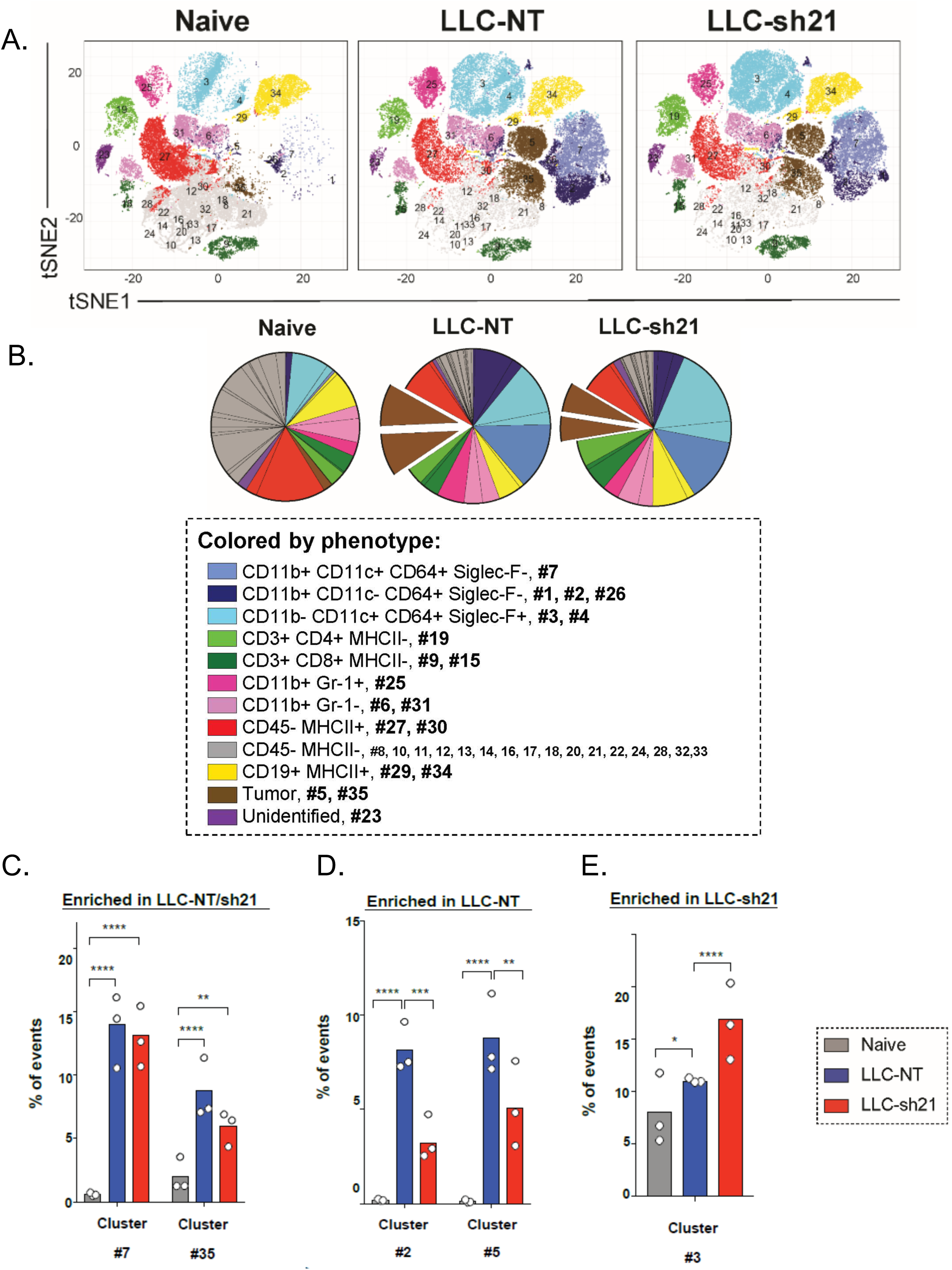
Socs1 KD Tumors Show Alterations in Multiple Populations. LLC-NT or LLC-sh21 (*Socs1* knockdown) cells were orthotopically injected into the left lung lobes of mice and established primary tumors. After 2 weeks of tumor growth with no treatment, mice were sacrificed and their tumor-bearing lung lobes were isolated. Single cell suspensions were made from tumor bearing lung lobes or naïve lungs. There were 3 experimental replicates each representing a pool of 3 tumors. Single cell suspensions of “Naïve”, “LLC-NT” or “LLC-sh21” were stained with a 39-antibody panel and analyzed on the Helios mass cytometer. Data show all viable single cells, subjected to the PhenoGraph algorithm. (A) PhenoGraph-defined cellular distribution and clustering, as defined by tSNE1 and tSNE2, colored by phenotypic designation (legend provided in panel B) for all treatment conditions where all replicates per experimental condition are combined. (B) Pie charts show all 35 clusters colored by their phenotypic designations for all experimental conditions with numbers indicating which PhenoGraph defined clusters were present in each phenotypic designation. Clusters identified as statistically significant are shown as preferentially enriched in: (C) Both the LLC-NT and LLC-sh21 tumor samples, (D) LLC-NT samples alone, or (E) LLC-sh21 samples alone. Error bars represent the mean of the data ±SEM after a two-way ANOVA (4C-E) (*p<0.05, **p<0.01, ***p<0.001, ****p<0.0001).

**Figure 4B** shows these data as percentages in pie graph format. Since cancer cells do not express a unique cell surface marker, we defined them as a CD45^-^ population that was absent in samples from naïve mice and highly enriched in the LLC-NT and LLC-sh21 tumor-containing samples (**Figure 4C**). Further examination of cancer cell clusters revealed two cancer cell clusters present in both LLC-NT and LLC-sh21 tumors that were defined by differential Ki67 expression (Cluster #5, Ki67^+^; Cluster #35, Ki67^-^) (**Supplemental Figure 3D**). Interestingly, we observed a reduction in the percentage of events that were cancer cells in LLC-sh21 compared to LLC-NT (17.54% LLC-NT to 11.07% LLC-sh21), with a decreased frequency of Ki67^+^ proliferating cancer cells in LLC-sh21 versus LLC-NT tumors (**Figure 4D**, **Supplemental Figure 3D**). Both LLC-NT and LLC-sh21 tumor cells expressed MHC Class I, but had low levels of MHC Class II expression, suggesting selective interactions with CD8^+^, but not CD4^+^ T cells (**Supplemental Figure 3E**). Interestingly, PD-L1 expression was not high on cancer cell clusters in either the LLC-NT or LLC-sh21 groups, indicating that although LLC-sh21 cells induce PD-L1 expression *in vitro* after IFNγ treatment (**Supplemental Figure 2C, 2H**), this is not reflected *in vivo* (**Supplemental Figure 3E**).

In addition to populations of tumor cells, we examined differences in inflammatory and immune cells between LLC-NT and LLC-sh21 tumors by examining CD45^+^ cell populations. We have previously profiled macrophage populations in LLC tumors using conventional flow cytometry [15]. We found that a population of recruited macrophages, defined as CD11b^+^/CD11c^+^/CD64^+^/SiglecF^-^ macrophages (Cluster #7), was enriched in tumor-bearing lungs (LLC-NT and LLC-sh21) relative to naïve lungs (**Figure 4C**). In addition, a population of recruited monocytes, defined as CD11b^+^/CD11c^-^/CD64^+^/SiglecF^-^ monocytes (Cluster #2), was selectively enriched in LLC-NT tumors relative to LLC-sh21 tumors or naïve lungs (**Figure 4D**). Conversely, a subset of alveolar macrophages defined as CD11b^-^/CD11c^+^/CD64^+^/SiglecF^+^ macrophages (Cluster #3), was enriched in LLC-sh21 tumors compared to LLC-NT or naïve lungs (**Figure 4E**). Although not enriched to a significant degree, we also detected increased CD8^+^ and CD4^+^ T cells in LLC-sh21 tumors versus LLC-NT tumors (**Figure 4A-B**).

Since we observed low levels of PD-L1 on both LLC-NT and LLC-sh21 tumor cells *in vivo*, we wanted to determine if expression of PD-L1 on non-tumor cells was altered in mice implanted with LLC-sh21 *Socs1* knockdown cancer cells [13]. While the overall frequency of CD64^+^ myeloid cells was comparable between LLC-NT and LLC-sh21 tumors, the relative distribution of macrophages expressing variable PD-L1 levels was altered (**Figure 5A-B**). Recruited monocytes (Cluster #2: CD11b^+^/CD11c^-^/CD64^+^/SiglecF^-^ cells), had low levels of PD-L1 expression and were less abundant in LLC-sh21 tumors compared to LLC-NT tumors (**Figure 5A-C)** [15]. Recruited macrophages (Cluster #7: CD11b^+^/CD11c^+^/CD64^+^/SiglecF^-^ cells), which express intermediate levels of PD-L1, were unchanged (**Figure 5A-B, 5D)**. Resident alveolar macrophages (Clusters #3 and #4: CD11b^-^/CD11c^+^/CD64^+^/SiglecF^+^ cells), which have the highest level of expression of PD-L1 relative to the other macrophage subsets, were increased in LLC-sh21 tumors (**Figure 5A-B, 5E**). These data collectively indicate that there is an increase in PD-L1^hi^ and a reciprocal decrease in PD-L1^lo^ macrophages in LLC-sh21 tumors, which may explain the efficacy of anti-PD-1 on these tumors.

**Figure 5.**
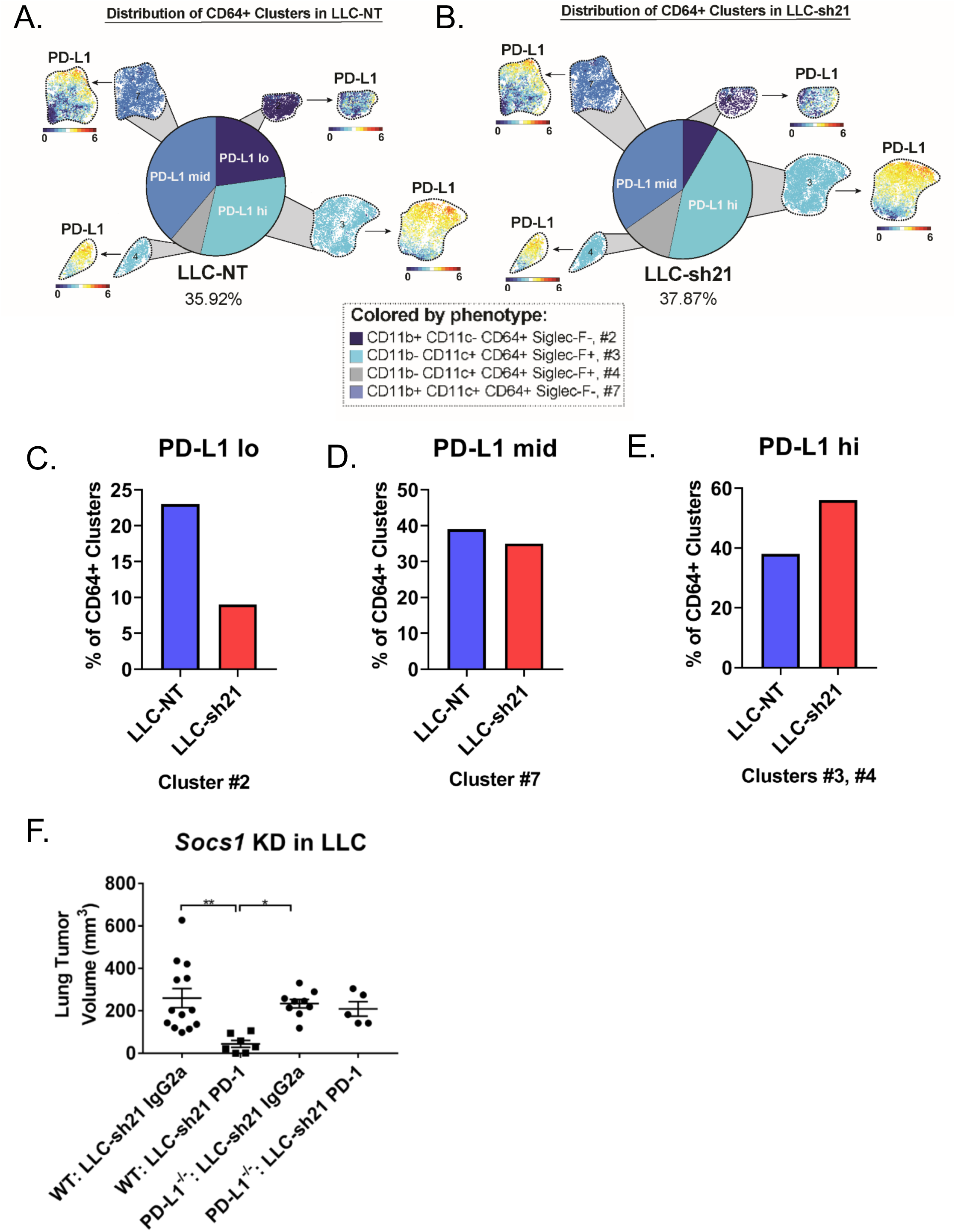
Socs1 KD Tumors Have An Altered Macrophage Composition. Pie charts show the relative frequency of cell clusters containing CD64^+^ events (a pan macrophage marker) in (A) LLC-NT or (B) LLC-sh21 tumors. Clusters are colored according to PD-L1 expression. Pie charts are quantified based on: (C) PD-L1 lo(Cluster #2), (D) PD-L1 mid(Cluster #7) or (E) PD-L1 hi(Clusters #3 and #4) expression.(F) LLC-sh21 cells were orthotopically injected into the lungs of either wild-type (WT) or PD-L1^-/-^ mice, established for 7 days, then were treated with either an isotype control antibody (IgG2a) or an anti-PD-1 antibody for 2 weeks followed by terminal sacrifice at 3 weeks post tumor cell injection. Primary tumor volume was assessed via digital calipers. Error bars represent the mean of the data ±SEM after a one-way ANOVA (5F)(*p<0.05, **p<0.01, ***p<0.001, ****p<0.0001).

In order to confirm the importance of PD-L1 expression on cells of the TME versus tumor-intrinsic PD-L1, we compared the response of LLC-sh21 tumors implanted into WT or PD-L1^-/-^ mice to anti-PD-1 treatment. When implanted into PD-L1^-/-^ mice, LLC-sh21 tumors lose their sensitivity to anti-PD-1 (**Figure 5F**). These results indicate that the PD-L1 from the tumor microenvironment is critical for host response to anti-PD-1 treatment.

### Socs1 KD Tumors Exhibit a More T-cell Inflamed Phenotype and Alterations in Macrophage Composition

While not statistically significant, we observed an increase in CD4^+^ and CD8^+^ T cell populations in LLC-sh21 tumors relative to LLC-NT tumors. Since T cells are critical for responses to immune checkpoint inhibitors, we further characterized changes in T cell populations by immunostaining. Representative images of both LLC-NT and LLC-sh21 tumors harvested at two weeks without treatment are shown (**Figure 6A-B, Supplemental Figure 4A-B**). Quantification of cells per high-power field (40X magnification) showed that LLC-sh21 tumors had significant increases in CD3^+^ T cells, a pan T cell marker, as well as trending increases in CD8^+^ and CD4^+^ populations relative to LLC-NT tumors (**Figure 6C-D, Supplemental Figure 4C**), consistent with our CyTOF data.

**Figure 6.**
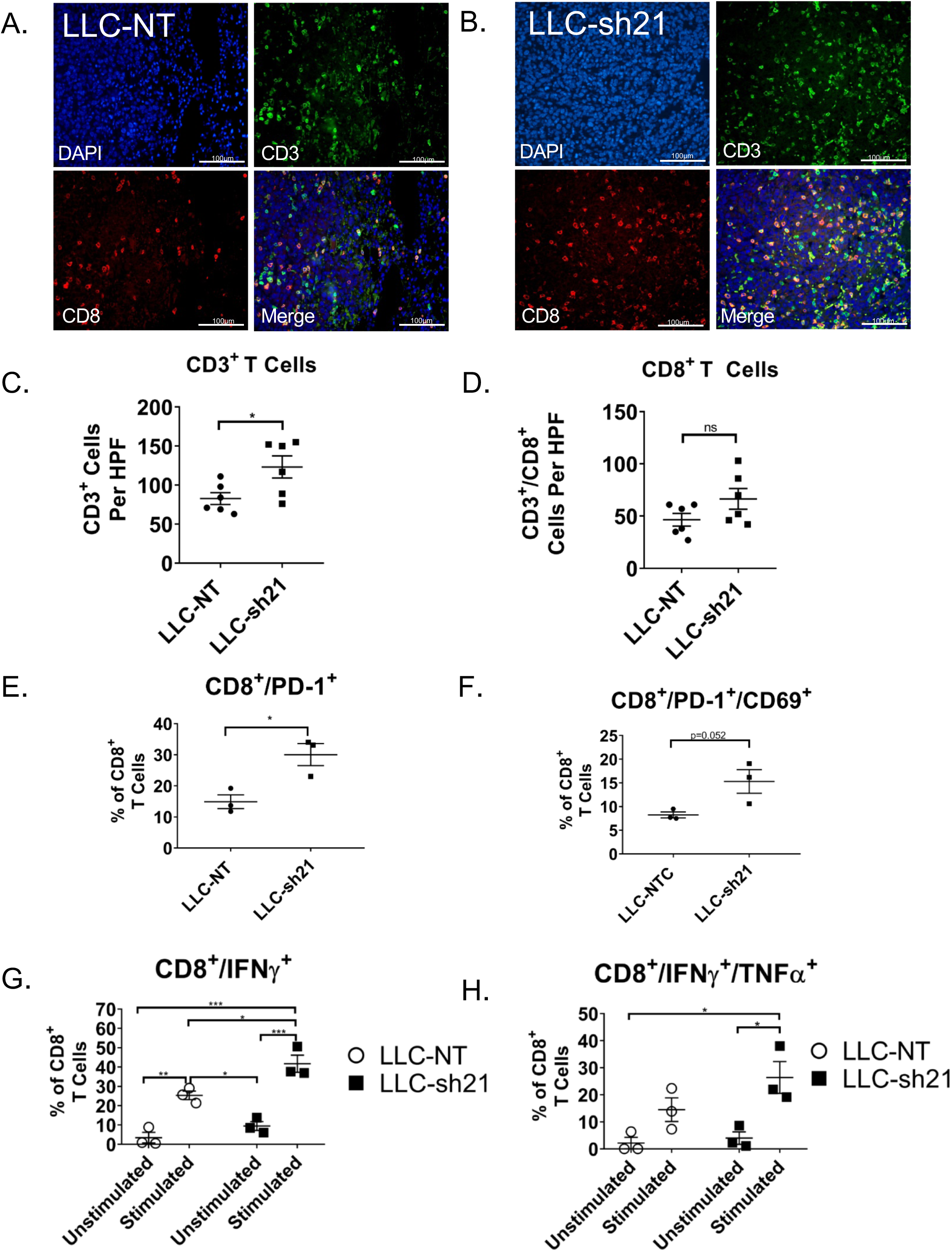
Socs1 KD Tumors Exhibit A More T Cell Inflamed Phenotype and Increased CD8^+^ T Cell Activation. LLC-NT or LLC-sh21 (*Socs1* knockdown) cells were injected into the left lung lobes of mice. After 2 weeks, mice were sacrificed and their tumor-bearing lung lobes were isolated for either flow cytometry or FFPE and T cell staining by immunofluorescence. Representative images of T cell staining from (A) A LLC-NT tumor or (B) A LLC-sh21 tumor at 40X magnification (Scale Bar 100µm) showing CD3^+^ cells (green), CD8^+^ cells, (red) and DAPI (blue). Quantification of (C) CD3^+^ T cells and (D) CD8^+^ T cells per high power field (HPF) in LLC-NT versus LLC-sh21 tumors (6 tumors in each group). T cell numbers per HPF were averaged over 4 experiments (6 random fields per tumor × 4 staining experiments=24 fields averaged/tumor in total). (E-H) Flow cytometry analysis of T cells was performed using 3 experimental replicates × 3 tumor bearing lung lobes each, for a total of 9 lung lobes per the experimental conditions of “LLC-NT” or “LLC-sh21”. (E) the percentage of PD-1 expressing CD8^+^ T cells, (F) double-positive PD-1/CD69 expressing CD8^+^ T cells gated as a percentage of all CD8^+^ T cells. (G-H) Single cell suspensions were either unstimulated (treated with Brefeldin and Monensin alone) or stimulated (treated with Brefeldin, Monensin, and PMA/Ionomycin) for 5 hours and analyzed by flow cytometry for (G) the percentage of single positive IFNγ expressing CD8^+^ T cells or (H) double-positive IFNγ/TNFα expressing CD8^+^ T cells gated as a percentage of all CD8^+^ T cells. Empty circles represent the LLC-NT condition while the solid black squares represent the LLC-sh21 condition. Error bars represent the mean of the data ±SEM after a student’s t-test (6C-F) or a two-way ANOVA (6G-H) (*p<0.05, **p<0.01, ***p<0.001, ****p<0.0001).

Changes in T cells were also analyzed by flow cytometry (**Supplemental Figure 4D**). By flow, we observed a significant increase in CD8^+^ T cells expressing PD-1 (**Figure 6E**) and a trending increase in the CD8^+^/PD-1^+^/CD69^+^ cells in LLC-sh21 versus LLC-NT tumors (**Figure 6F**), indicating that a significant percentage of PD-1 expressing CD8^+^ T cells had recently been exposed to antigen. These changes were not observed in CD4^+^ T cells **(Supplemental Figure 4E-F**) Upon cellular stimulation with PMA/Ionomycin in the presence of Golgi inhibitors, we detected significant increases in IFNγ-positive and IFNγ/ TNFα double-positive CD8^+^ T cells in LLC-sh21 tumors versus LLC-NT tumors (**Figure 6G-H**), indicating more cytotoxic and anti-tumor capacities. Similar increases were observed in CD4+ T cells though these were not statistically significant (**Supplemental Figure 4G-H**). These data indicate that LLC-sh21 tumors at baseline, prior to anti-PD-1 therapy, have greater CD8^+^ T cell activation and by extension, recognition of tumor cells than LLC-NT tumors.

### Socs1 KD Tumors Have Elevated Levels of Cxcl9

Since we observed increased tumor-infiltrating T cells in LLC-sh21 *Socs1* KD tumors, as well as increased CD8^+^ T cell activation, we sought to identify cancer cell-intrinsic factors that could mediate these effects. We therefore recovered LLC-NT and LLC-sh21 cancer cells implanted into GFP-expressing transgenic mice and compared gene expression profiles with the respective cells grown *in vitro* using RNA-Seq (as in **Figure 1**). These data showed that *Socs1* expression was decreased by approximately 60% in LLC-sh21 compared LLC-NT tumors, confirming that these cells *in vivo* were still silenced for *Socs1* (data not shown). We identified 45 genes that were differentially expressed between the LLC-NT and LLC-sh21 cells *in vivo* (**Supplemental Table 1**) that met our criteria of q<0.03. Of interest, expression of *Cxcl9* as well as three MHC Class I genes (*H2k1*, *H2q1*, *H2q4*) was increased in LLC-sh21 cells *in vivo* compared to the LLC-NT cells (**Figure 7A**, **Supplemental Table 1**). We confirmed changes in *Cxcl9* by in situ hybridization. Relative to tumor sections stained with a negative control dapB (**Figure 7B**), or a normal adjacent lung stained with a probe for murine *Cxcl9* (another negative control) (**Figure 7C**), 4 separate LLC-NT or LLC-sh21 tumors stained positive for *Cxcl9* (**Figure 7D-E**). There was much higher *Cxcl9* staining in 3 out of 4 LLC-sh21 tumors, particularly around the tumor edge compared to LLC-NT tumors. While we do not exclude other potential tumor-intrinsic mechanisms, the increased levels of *Cxcl9* within and around LLC-sh21 tumors would allow for increased T cell infiltration and trafficking into tumors.

**Figure 7.**
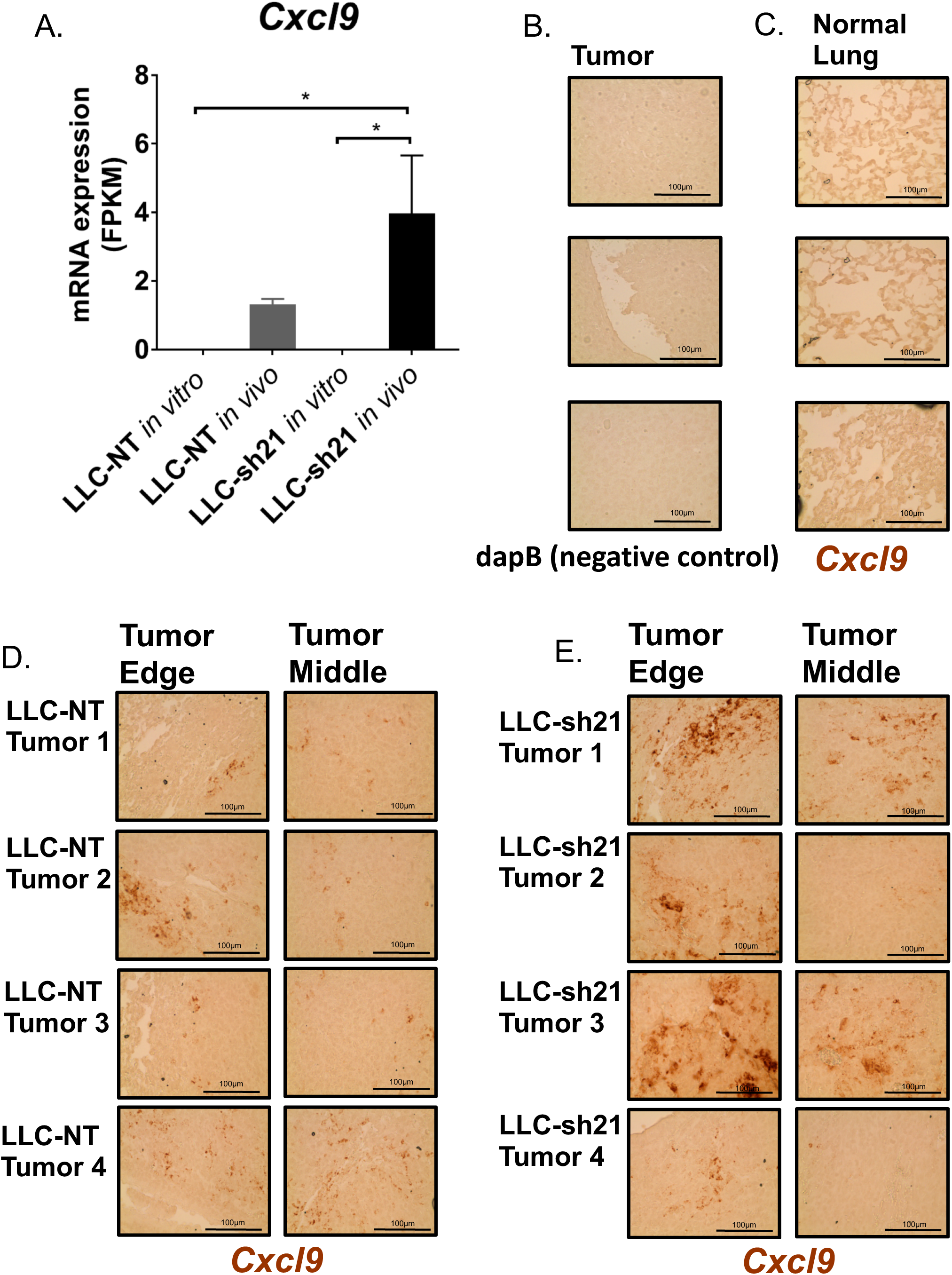
Socs1 KD Tumors Have Elevated Levels of Cxcl9. (A) LLC-NT or LLC-sh21 (*Socs1* knockdown) cells were injected into the left lung lobe of transgenic GFP-expressing C57BL/6J mice and were grown for 3 weeks. Single cell suspensions underwent FACS analysis as in Figure 1, and RNA was isolated from GFP-negative (tumor cells). RNA-Seq was performed and compared with RNA isolated from identical cells grown *in vitro*. Both the LLC-NT and LLC-sh21 conditions had 3 experimental replicates per *in vitro* and *in vivo* conditions with five tumor-bearing lung lobes pooled per *in vivo* experimental replicate (15 mice used total). *In vitro and in vivo* mRNA expression of *Cxcl9* in FPKM as assessed by RNA-Seq. (B-D) The RNAScope system was used for In Situ Hybridization. Pictures were taken at 40X (Scale Bar 100µm) (B) Tumor tissue stained with a negative control probe, dapB. (C) Adjacent normal lung stained with a probe targeting murine *Cxcl9* that shows negative staining. (D) 4 separate tumors from either the LLC-NT or (E) LLC-sh21 experimental conditions stained with a probe targeting murine *Cxcl9*. Dark brown regions represent positive Cxcl9 staining. Error bars represent the mean of the data ±SEM after a one-way ANOVA (7A). (*p<0.05, **p<0.01, ***p<0.001, ****p<0.0001).

## Discussion

While biomarkers have been developed which correlate the response of lung cancer patients to anti-PD1/anti-PD-L1 therapy, defining the cellular and molecular pathways that regulate this response remains poorly understood. We have previously demonstrated differential responses to anti-PD-1/anti-PD-L1 of two K-Ras mutant lung cancer cells, with CMT167 tumors showing a strong inhibition, and LLC tumors being resistant to therapy [13]. In this study we have sought to define intrinsic features of the cancer cell that mediate this differential response. Our data define responsiveness of cancer cells to IFNγ as a critical determinant to responsiveness to anti-PD-1 therapy. Furthermore, we have shown that altering responsiveness of the cancer cells to IFNγ causes complex multifaceted changes in the microenvironment.

By analyzing gene expression changes *in vivo*, we determined that LLC cells failed to robustly induce an IFNγ signature compared to CMT167 tumors. An IFNγ signature in bulk tumor tissue (which includes a complex mixture of cancer cells, stromal cells, and hematopoietic cells, etc.) has been associated with responsiveness to anti-PD-1 therapy in lung cancer and others malignancies [9]. However, whether this is associated with cancer cells alone or the surrounding TME has not been well examined. We hypothesized that sensitivity of cancer cells to IFNγ is a major regulator of the TME, and that altering the sensitivity of the cancer cells would regulate the response of tumors to checkpoint inhibition. In our studies we have selectively altered expression of a single gene in these cells, and therefore not affected their mutational burden.

In CMT167 cells, silencing the expression of *Ifngr1* decreased JAK/STAT signaling and made tumors resistant to anti-PD-1 therapy. Similar results have been obtained in melanoma models [30]. This effect does not appear to be a consequence of altered proliferation of the silenced cells, but was associated with decreased infiltration of T cells into the tumors, consistent with the model proposed that less-inflamed tumors are resistant to immunotherapy [5, 31]. Conversely, our data show that LLC cells have a diminished response to IFNγ *in vitro*, associated with high basal expression of *Socs1*, and silencing *Socs1* markedly increases the response to IFNγ *in vitro* through enhanced JAK/STAT signaling. Importantly, cancer cells with *Socs1* silencing (LLC-sh21) grown as orthotopic tumors are much more responsive to anti-PD-1 therapy than LLC-NT controls. This is not observed if the identical cells are implanted subcutaneously, suggesting that silencing *Socs1* alters communication between cancer cells and specific components of the lung microenvironment that are not present in subcutaneous tumors (e.g. resident alveolar macrophages). Several previous studies have identified *Socs1* as a gene associated with immunosuppression and tumor progression. In melanoma, increased copy number of *Socs1* was found in tumors resistant to anti-CTLA therapy[30]. In human lung tumors, MET activation was associated with increased *Socs1* expression and escape from immunotherapy [32]. Conversely, chemotherapeutic agents were shown to down regulate *Socs1* through induction of *miR-155*, resulting in increased activation of CD8^+^ T cells [33].

We anticipated that LLC-sh21 cells would express higher levels of CXCL9 *in vivo*, which we detected by message in tumor sections. Due to elevated *Cxcl9* expression in these tumors, there were more recruited and activated T cells--particularly CD8^+^ T cells. These results are indicative of a more “T cell inflamed tumor” which is associated with better response to checkpoint inhibitors [5]. Though we anticipated increased expression of PD-L1 on LLC-sh21 cancer cells *in vivo*, which would promote a T cell-rich microenvironment with high PD-L1 expression, and subsequent sensitivity to anti-PD-1, LLC-sh21 cells recovered from tumors only showed a modest induction of PD-L1 compared to LLC-NT cells (data not shown). This could be explained by only a subset of the cancer cells expressing PD-L1, for example at the tumor edge. In our previous studies, PD-L1 expression on CMT167 cells, which are sensitive to IFNγ and checkpoint inhibition, was not detected on all cancer cells [13].

Unexpectedly, our data indicate that LLC-sh21 tumors also exhibit tumor extrinsic changes compared to LLC-NT tumors of equal size. We observed a decrease in the percentage of cancer cells in LLC-sh21 compared to LLC-NT, and a reciprocal increase in T cells, which confirmed our immunostaining data. In addition, we observed complex changes in in the myeloid compartment, with proportional increases in resident alveolar macrophages and decreases in recruited monocytes/macrophages. Importantly, the resident alveolar population expressed high levels of PD-L1, whereas the populations that decreased had moderate to low expression of PD-L1. Thus, silencing of *Socs1* in the cancer cells results in a tumor with increased numbers of PD-L1^hi^ macrophages. Again, our previous study demonstrated that a critical difference between CMT167 and LLC tumors was markedly increased numbers of PD-L1^hi^ macrophages on CMT167 tumors [13]. Therefore, silencing *Socs1* has converted LLC tumors to have many of the features of CMT167 tumors: increased T cell recruitment, increased levels of CXCL9 and increased numbers of PD-L1^hi^ macrophages. Decreased levels of *Ccl2* in LLC-sh21 tumors, demonstrated by our RNA-Seq data (**Supplemental Table 1**) may account for the decreased frequencies of recruited macrophage/monocyte populations— potentially immunosuppressive populations [15]. The role of this factor requires further examination in this model. Consistent with a important role for lung-specific macrophage populations,, we propose that the lack of alveolar macrophages (PD-L1^hi^) in the subcutaneous model are responsible for the lack of response of subcutaneous LLC-sh21 tumors to anti-PD-1 therapy.

In summary, our data identify a critical role for IFNγ sensitivity within cancer cells as a major determinant that directly shapes the TME. The results also underscore the complex interplay between cancer cells and populations of inflammatory and immune cells. These interactions are mediated through production of paracrine factors, including chemokines and potentially lipid mediators. Subtle changes, such as altering expression of a single gene in the cancer cells changes these interactions in profound ways, likely by altering the secretome of the cancer cells. Therapeutically, in this case, we have generated a tumor with increased T cells and fewer myeloid cells, which is associated with an increased response to anti-PD-1. However, the complexity of the crosstalk suggests that a better understanding of how the various cell populations interact is needed to design more effective combination therapies for treatment of lung cancer.

## Acknowledgements

We thank Lynn Heasley for helpful discussion.

## Notes

**Financial Support:** This work was supported by the NIH (R01 CA162226 and CA236222 to RA Nemenoff), Colorado Lung SPORE (P50 CA058187 to HY Lee and RA Nemenoff) and the United States Department of Veterans Affairs Biomedical Laboratory Research and Development Service (Career Development Award IK2BX001282 to HY Li). The University of Colorado Cancer Center Flow Cytometry and the Genomics and Microarray Shared Resources is supported by NIH P30CA046934. The University of Colorado Cancer Center Flow Cytometry Core Facility is funded through a support grant from the National Cancer Institute (P30 CA046934). Imaging experiments were performed in the University of Colorado Anschutz Medical Campus Advanced Light Microscopy Core supported in part by NIH/NCATS Colorado CTSI Grant Number UL1 TR001082. None of the authors have any conflicts of interest.

