## Supplemental Figures 1-5 and Table for "Tumor-Intrinsic Response to IFNγ Shapes the Tumor Microenvironment and Anti-PD-1 Response in NSCLC"

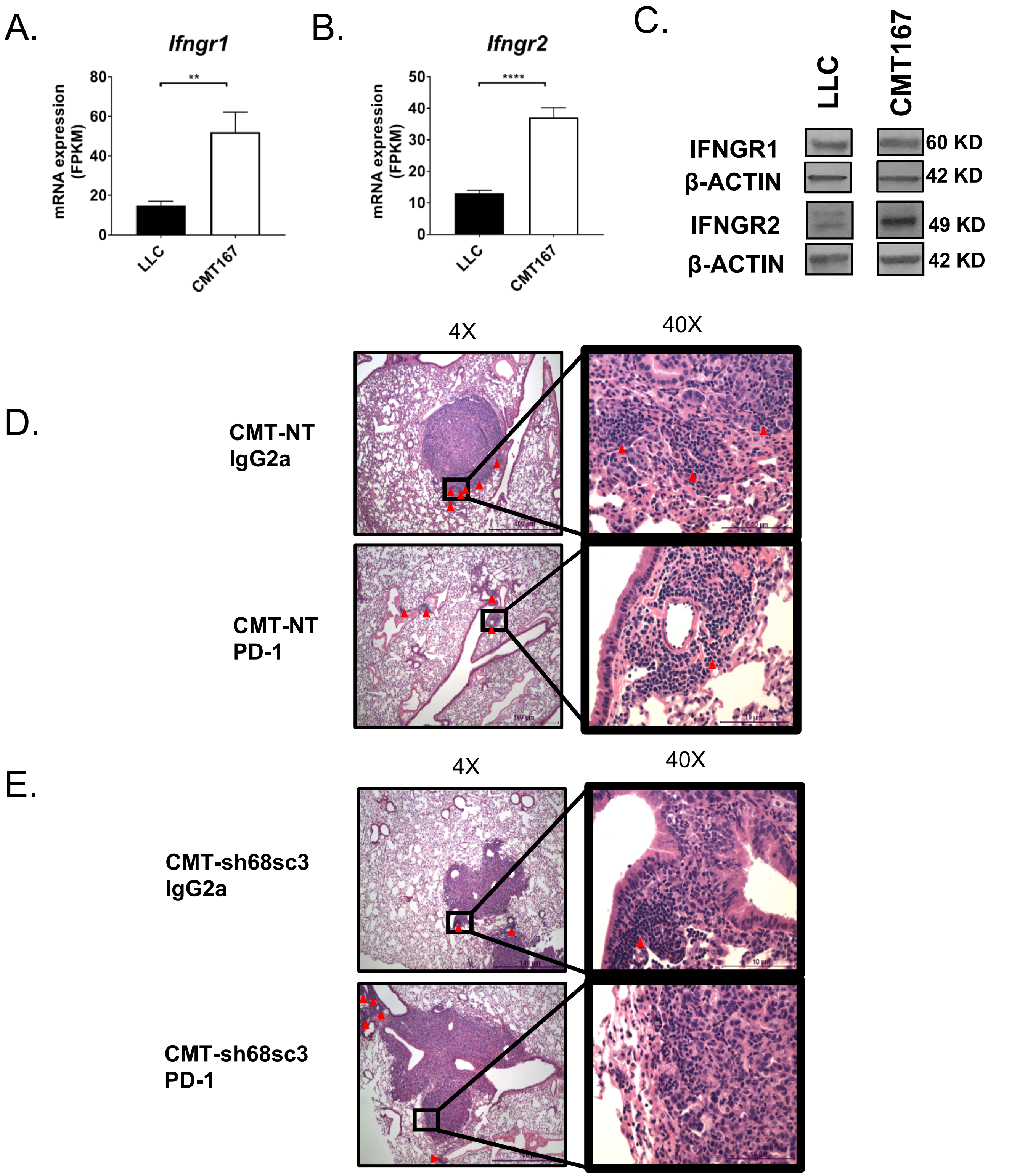


**Supplemental Figure 1**. *In vitro* mRNA expression of *Ifngr1* (A) and *Ifngr2* (B) in FPKM as assessed by RNA-Seq. (C) Immunoblots of IFNGR1 and IFNGR2 expression relative to β-ACTIN levels in LLC and CMT167 cells *in vitro*. Representative images of H&E staining of tumors from the following experimental conditions: CMT-NT IgG2a and CMT-NT PD-1 (D), and CMT-sh68sc3 IgG2a and CMT-sh68sc3 PD-1 (E) at 4X (Scale Bar 100μm) and 40X (Scale Bar 10μm) magnification. Tumors are dense and stain dark purple relative to adjacent normal lung tissue. Red arrows mark lymphocyte nests. Error bars represent the mean of the data ±SEM after a student's t-test (SF1A-B) (*p<0.05, **p<0.01, ***p<0.001, ****p<0.0001).

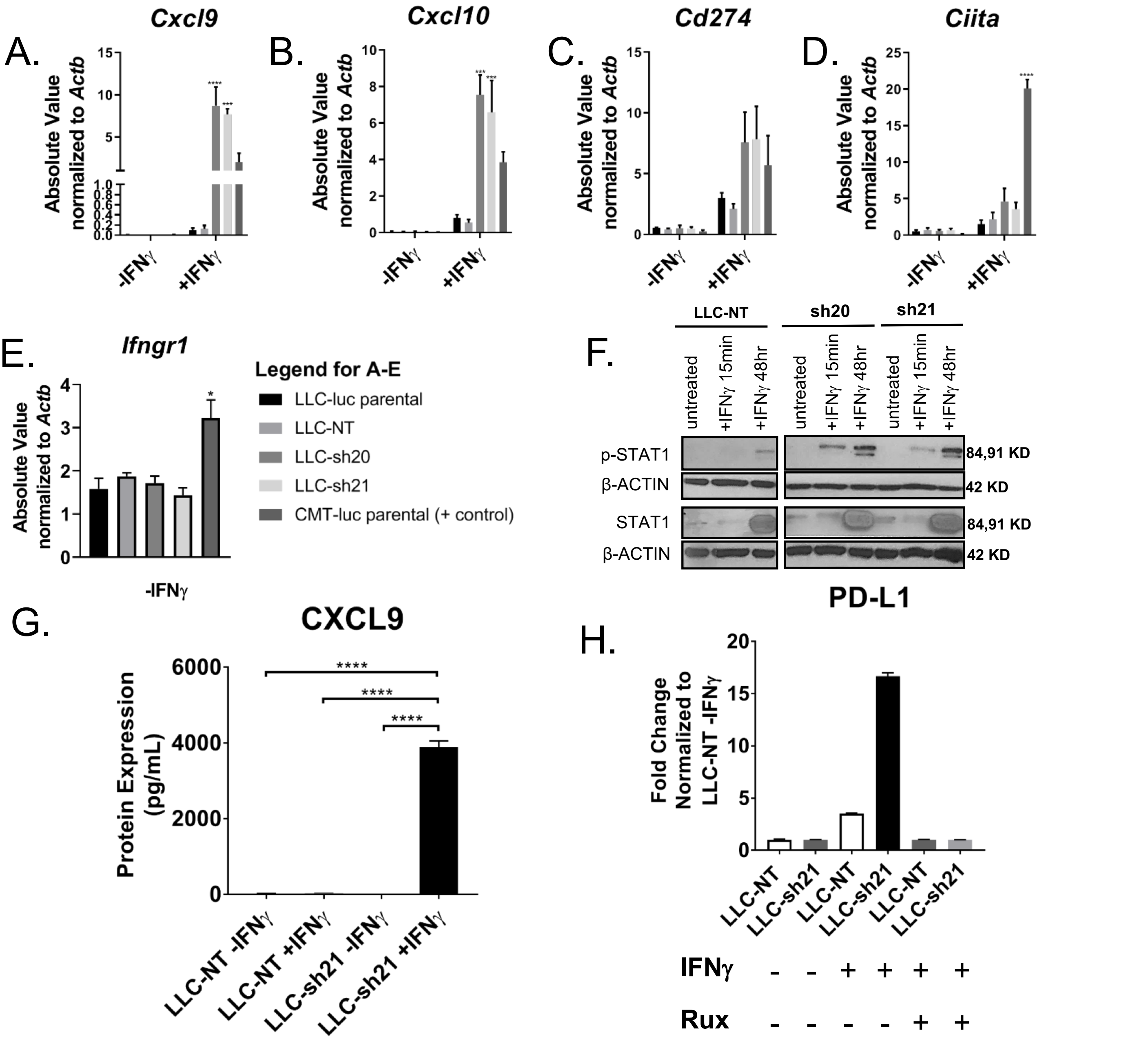


**Supplemental Figure 2**. 2 separate shRNAs targeting *Socs1* (LLC-sh20, LLC-sh21) and a non-targeting control vector (LLC-NT) were transduced into LLC cells expressing luciferase. LLC cells were then screened for functional and stable knockdown of *Socs1* after 10 days of puromycin selection. Cells were treated with ± 10ng/mL IFNγ for 24 hours followed by isolation of RNA and qRT-PCR. mRNA levels of (A) *Cxcl9*,(B) *Cxcl10*, (C) *Cd274*, (D) *Ciita* and (E) *Ifngr1* are shown as Absolute Values (SQ Values) normalized to the housekeeping gene *Actb*. Statistics compare the CMT-NT line to other cell lines with or without treatment. (F) Immunoblots showing p-STAT1, total STAT1, and β-ACTIN levels of the CMT-NT, CMT-sh68sc3, and CMT-sh69sc2 cell lines ±IFNγ after 15 minutes or 48 hours *in vitro*. (G) Conditioned media from cells plated at equal confluency was measured for CXCL9 protein via ELISA comparing the LLC-NT line to LLC-sh21 cells *in vitro*. Error bars represent the mean of the data ±SEM (*p<0.05, **p<0.01, ***p<0.001, ****p<0.0001). (H) PD-L1 protein expression *in vitro* as measured by flow cytometry ± IFNγ for 18 hours and ± 1μM Ruxolitinib (JAK1/JAK2 inhibitor) or vehicle control. Data was normalized to the median fluorescence intensity (MFI) of the LLC-NT -IFNγ sample. All bars are ****p<0.0001 except untreated cells compared to +IFNγ+Rux cells. A two way ANOVA was used to analyze SF2A-D, and a one-way ANOVA was used to analyze SF1E, and G-H.

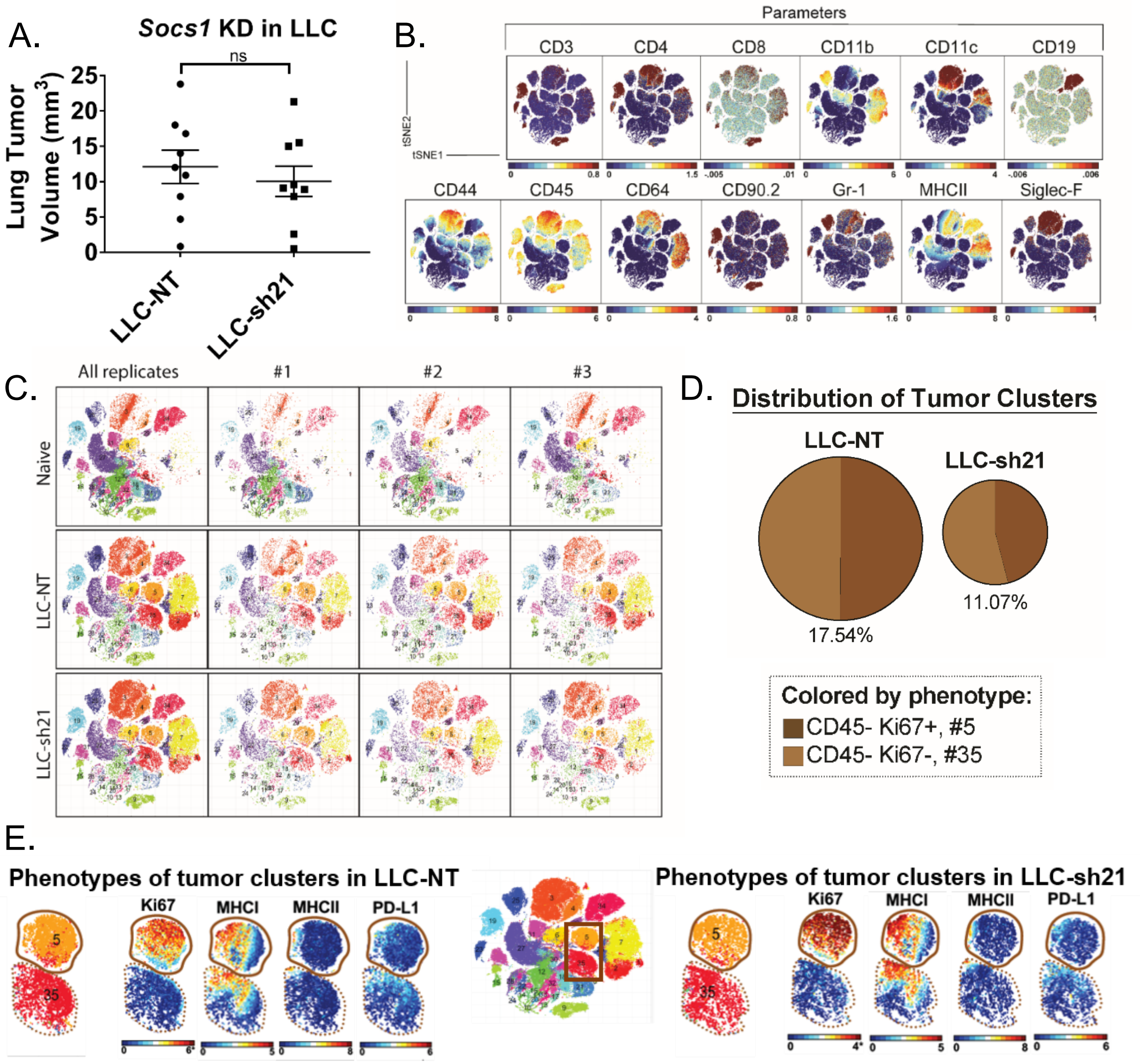


**Supplemental Figure 3.** LLC-NT or LLC-sh21 (Socs1 knockdown) cells were orthotopically injected into the left lung lobes of mice and established primary tumors. After 2 weeks of tumor growth with no treatment, mice were sacrificed and their tumor-bearing lung lobes were isolated. Single cell suspensions were made from tumor bearing lung lobes or naïve lungs. There were 3 experimental replicates x 3 tumor-bearing lung lobes each, for a total of 9 lung lobes per the experimental conditions of “Naïve”, “LLC-NT” or “LLC-sh21”. Single cell suspensions were then stained with a 39-antibody panel and analyzed on the Helios mass cytometer. Data show all viable single cells, subjected to the PhenoGraph algorithm. (A) Primary tumor volume was assessed by digital calipers. (B) PhenoGraph-based visualization on a tSNE plot, colored according to expression of the cellular markers (parameters): CD3, CD4, CD8, CD11b, CD11c, CD19, CD44, CD45, CD64, CD90.2, Gr-1, MHC-II, and Siglec-F. (C) PhenoGraph-defined cellular distribution and clustering, as defined by tSNE1 and tSNE2, colored by cluster ID for all experimental groups and replicates.(D) Pie charts show the abundance of all clusters containing tumor cells (identified as $CD45^{-}$ and missing in naïve samples). (E) Tumor cell clusters in LLC-NT samples or LLC-sh21 samples colored according to Ki67, MHCI, MHCII, and PD-L1.

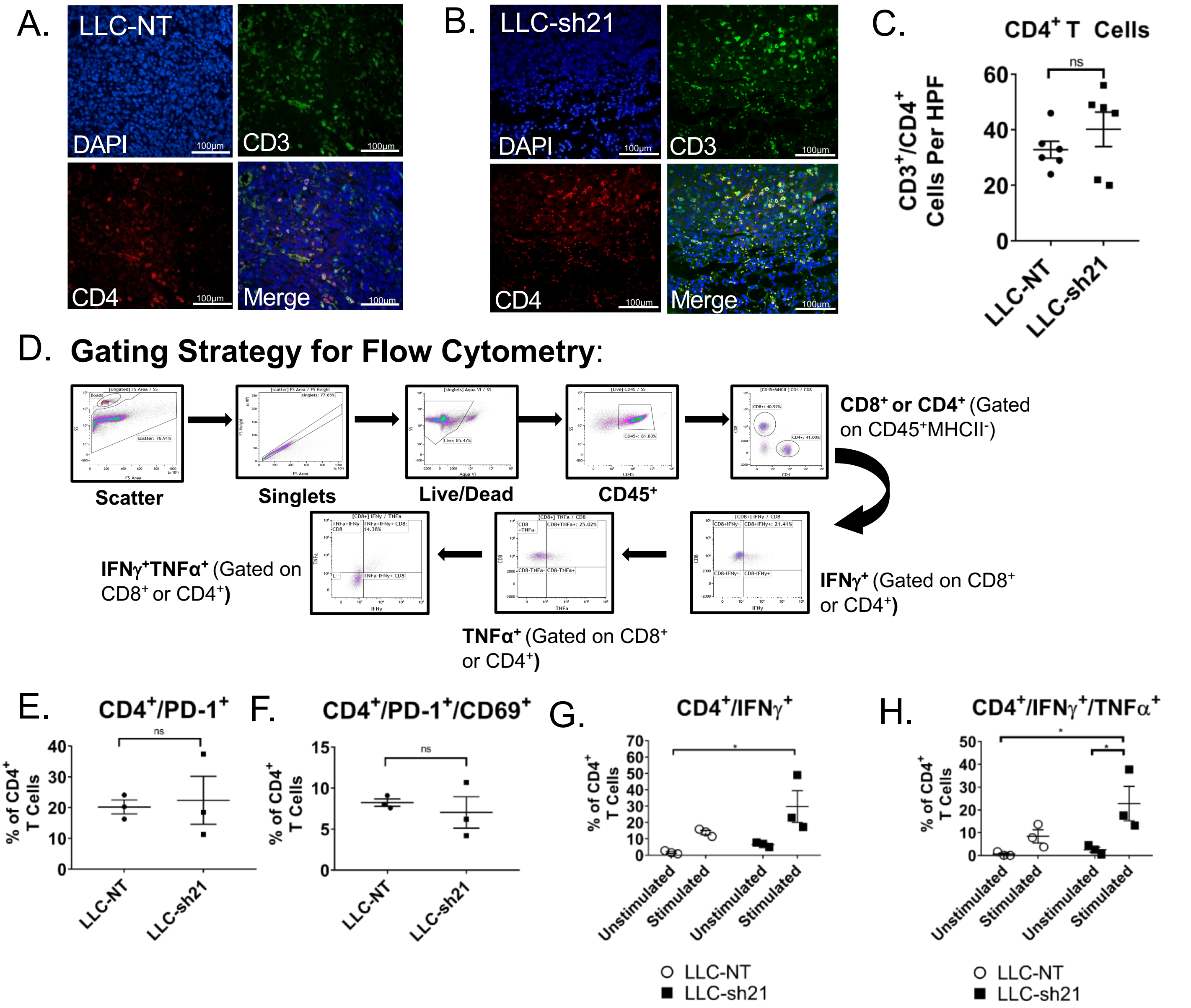


**Supplemental Figure 4**. LLC-NT or LLC-sh21 (Socs1 knockdown) cells were orthotopically injected into the left lung lobes of mice and established primary tumors. After 2 weeks of tumor growth with no treatment, mice were sacrificed and their tumor-bearing lung lobes were isolated for either flow cytometry or FFPE and T cell staining by immunofluorescence. Representative images of T cell staining from an (A) a LLC-NT tumor or (B) a LLC-sh21 tumor at 40X magnification (Scale Bar 100µm) showing CD4+ T cell staining. DAPI is shown in blue, CD3 in green, CD4 in red, and a Merge of all channels in yellow. (C) Quantification of CD4+ T cells and per high power field (HPF) in LLC-NT versus LLC-sh21 tumors. (D) Gating Strategy for Flow Cytometry (**Figure 6**). Gates Determined Based on Isotype Controls (for IFNγ and TNFα) or Single Stain Controls for Aqua Viability, CD45, MHCII, CD8, or CD4. (E) Percentage of PD-1 expressing CD4+ T cells or (F) double-positive PD-1/CD69 expressing CD4+ T cells gated as a percentage of all CD4+ T cells. (G) Percentage of single positive IFNγ expressing CD4+ T cells or (H) double-positive IFNγ/TNFα expressing CD4+ T cells gated as a percentage of all CD4+ T cells. For G-H, empty circles represent the LLC-NT condition while the solid black squares represent the LLC-sh21 condition.  Error bars represent the mean of the data ±SEM after a student's t-test (SF4E-F) or a two-way ANOVA (SF4G-H) (*p<0.05, **p<0.01, ***p<0.001, ****p<0.0001).

| Column1 | LLC-NT *in vivo* Mean | LLC-sh21 *in vivo* Mean | q value | significance |
|---|---|---|---|---|
| Acpp | 12.1419 | 6.39922 | 0.0172688 | yes |
| Acsbg1 | 5.82487 | 2.91786 | 0.0470298 | yes |
| Acta2 | 3.92554 | 7.91805 | 0.0172688 | yes |
| Adam19 | 0.240938 | 0.963345 | 0.0172688 | yes |
| Adamts1 | 41.17 | 63.8169 | 0.0172688 | yes |
| Adamts14 | 13.3338 | 8.91634 | 0.0290842 | yes |
| Add3 | 2.45631 | 4.56274 | 0.0172688 | yes |
| Arl4a | 21.6384 | 33.0467 | 0.0290842 | yes |
| Ccbe1 | 85.8788 | 60.3206 | 0.0172688 | yes |
| Ccl2 | 330.633 | 224.784 | 0.0404341 | yes |
| Cfhr2 | 10.0291 | 21.6197 | 0.0172687 | yes |
| Clip3 | 4.08066 | 7.74605 | 0.0172688 | yes |
| Col1a1 | 49.5513 | 79.3589 | 0.0172688 | yes |
| Colec12 | 20.5231 | 30.2074 | 0.0290842 | yes |
| Cxcl9 | 1.31311 | 3.96966 | 0.0172688 | yes |
| Ece1 | 2.00138 | 3.96191 | 0.0172688 | yes |
| Edil3 | 19.7025 | 34.3739 | 0.0172688 | yes |
| Eya1 | 18.3614 | 27.6141 | 0.0172688 | yes |
| Fam167a | 18.5372 | 11.5549 | 0.0172688 | yes |
| Fgf10 | 3.88817 | 6.98255 | 0.0172688 | yes |
| Foxc1 | 6.78145 | 12.3038 | 0.0172688 | yes |
| Frzb | 1.2358 | 3.74932 | 0.0172688 | yes |
| H2-K1 | 37.5817 | 63.8102 | 0.0172688 | yes |
| H2-Q1 | 59.0791 | 89.1811 | 0.0290842 | yes |
| H2-Q4 | 42.5519 | 64.8862 | 0.0172688 | yes |
| Il18rap | 41.0198 | 27.0563 | 0.0172688 | yes |
| Mfap4 | 42.4817 | 77.9126 | 0.0172688 | yes |
| Mndal | 12.2216 | 18.5332 | 0.0404341 | yes |
| Nod1 | 0.697882 | 2.08266 | 0.0290842 | yes |
| Nov | 23.5182 | 51.0226 | 0.0172688 | yes |
| Olr1 | 4.23748 | 1.56871 | 0.0172688 | yes |
| P4ha3 | 5.6243 | 8.99315 | 0.0470298 | yes |
| Palld | 4.69332 | 7.67283 | 0.0470298 | yes |
| Pf4 | 5.69022 | 0 | 0.0172688 | yes |
| Pgf | 0.764998 | 2.63044 | 0.0470298 | yes |
| Rassf9 | 1.87097 | 5.64848 | 0.0172688 | yes |
| Rcn3 | 1.32811 | 5.04325 | 0.0172688 | yes |
| Sema7a | 12.2671 | 6.61693 | 0.0172688 | yes |
| Selenop | 22.0343 | 35.9424 | 0.0172688 | yes |
| Thsd7a | 0.550461 | 1.14069 | 0.0470298 | yes |
| Tppp3 | 31.2157 | 49.0393 | 0.0404341 | yes |
| Ubc | 5.81277 | 10.454 | 0.0172688 | yes |
| Xlr5c | 1.44416 | 0 | 0.0172688 | yes |
| Zfp820 | 2.0774 | 4.01065 | 0.0290842 | yes |

**Supplemental Table 1.** LLC-NT or LLC-sh21 (*Socs1* knockdown) cells were orthotopically injected into the left lung lobe of transgenic GFP-expressing C57BL/6J mice and were grown for 3 weeks. Tumor-bearing lung lobes were isolated and made into single cell suspensions containing both GFP-positive (host cells) and GFP-negative (tumor cells). First, RNA was isolated from identical cells grown in passage (*in vitro* condition). Second, RNA was isolated from recovered GFP-negative tumor cells (isolated via FACS-*in vivo* condition). Third, RNA was run for RNA-Seq from both conditions. Both the LLC-NT and LLC-sh21 conditions had 3 experimental replicates per *in vitro* and *in vivo* conditions with five tumor-bearing lung lobes pooled per *in vivo* experimental replicate (15 mice used total). Table showing the top 45 differentially expressed genes between the *in vivo* RNA-Seq conditions in LLC-NT or LLC-sh21 tumors. Genes that are known to have important roles in the TME are highlighted in red.
